# Interactions of outer membrane lipoproteins *P. aeruginosa* PA3214 and *E. coli* PqiC with their MCE protein binding partners, PA3213 and PqiB

**DOI:** 10.64898/2026.05.09.724024

**Authors:** Sabrina I. Giacometti, Nicolas Coudray, Rachel L. Redler, Gira Bhabha, Damian C. Ekiert

**Affiliations:** Department of Biology, Johns Hopkins University, Baltimore, MD, USA; Applied Bioinformatics Laboratories, New York University School of Medicine, New York, USA; Structural Biochemistry Group; Department of Biology, Chemistry, and Pharmacy; Freie Universität Berlin

**Author notes:** For correspondence (GB) and (DCE).

## Abstract

Members of the Mammalian Cell Entry (MCE) superfamily interact with other proteins to form diverse architectures for the transport of hydrophobic molecules across the cell envelope in Gram-negative bacteria. Some of these trans-envelope MCE protein complexes include a PqiC-like outer membrane (OM) lipoprotein component. The best-studied member of this group of OM lipoproteins is *E. coli* PqiC, from the PqiABC system, which can form an octameric ring. How PqiC-like lipoproteins interact with their MCE protein binding partners to facilitate transport is not well understood. Here we report the cryo-electron microscopy structures of *Pseudomonas aeruginosa* PA3214, a homolog of PqiC, in the context of the full MCE transport PA3211-PA3214 system. Our structure provides insight into the biological assembly of the lipoprotein and interactions with its binding partner, MCE protein PA3213. We utilize deep mutational scanning to identify functionally important sites in *E. coli* PqiC in an unbiased manner. Through phenotypic and biochemical experiments, we characterize the interactions of the lipoproteins PqiC and PA3214 with their associated MCE proteins PqiB and PA3213, thus providing a model for how some MCE proteins employ a C-terminal peptide to mediate key interactions with their cognate lipoproteins at the OM.

## Introduction

The cell envelope of Gram-negative bacteria consists of an inner membrane (IM) and an outer membrane (OM) separated by an aqueous periplasmic space where the peptidoglycan resides. The OM is typically an asymmetric lipid bilayer containing lipopolysaccharides (LPS) in the outer leaflet and phospholipids in the inner leaflet (1). The OM forms a protective barrier surrounding the cell, but must also allow flux of specific molecules into and out of the cell. As such, specialized protein-based systems have evolved to facilitate: 1) Nutrient import into the cell; 2) Export of toxic molecules out of the cell, such as antibiotics; 3) Transport of lipids between the IM and OM to maintain OM integrity. Transport systems responsible for trans-envelope movement of hydrophobic molecules such as lipids are often driven by ATP hydrolysis in the cytoplasm (2–5), or utilize energy derived from the proton motive force (6– 8), or alternatively, may be passive transporters (9–12).

Mammalian Cell Entry (MCE) domain-containing proteins (MCE proteins) interact with other proteins to form macromolecular machines that transport phospholipids, cholesterol, and other hydrophobic molecules (13–22). MCE proteins are found in most double-membrane bacteria, as well as in plant chloroplasts (23, 24). Typically MCE systems consist of at least 2 main parts: 1) an inner membrane complex, which may include an ABC (ATP-Binding Cassette) transporter (13, 14, 16, 18–20, 25) or a LetA-like membrane protein (17), and 2) a periplasmic MCE protein, which are anchored to the IM by a single-pass transmembrane helix or, occasionally, by a lipid anchor (16, 26). Some MCE systems additionally contain a lipoprotein, thought to be anchored to the OM (**Fig. 1A**). MCE proteins oligomerize to form homo-(13, 15) or hetero-(16) hexameric rings, with a central pore through which substrates can pass. The overall architecture of the MCE proteins in structurally characterized systems, and correspondingly their mechanism of transport, is quite different (**Fig. S1**). While our functional understanding of the IM and MCE components of MCE systems in lipid transport has advanced greatly in recent years (14–18, 20, 25, 27–33), how lipids are inserted/extracted at the OM remains more obscure (34–37). In several MCE systems, such as *E. coli* Let (15, 38) and mycobacterial Mce1 (16), no clear OM components have been identified. In MCE systems with known OM-associated binding partners, the protein takes the form of a porin-like beta-barrel (37, 39) or a lipoprotein, which can adopt at least two distinct folds and includes both MlaA-like (34, 40) and PqiC-like proteins (41, 42).

**Figure 1.**
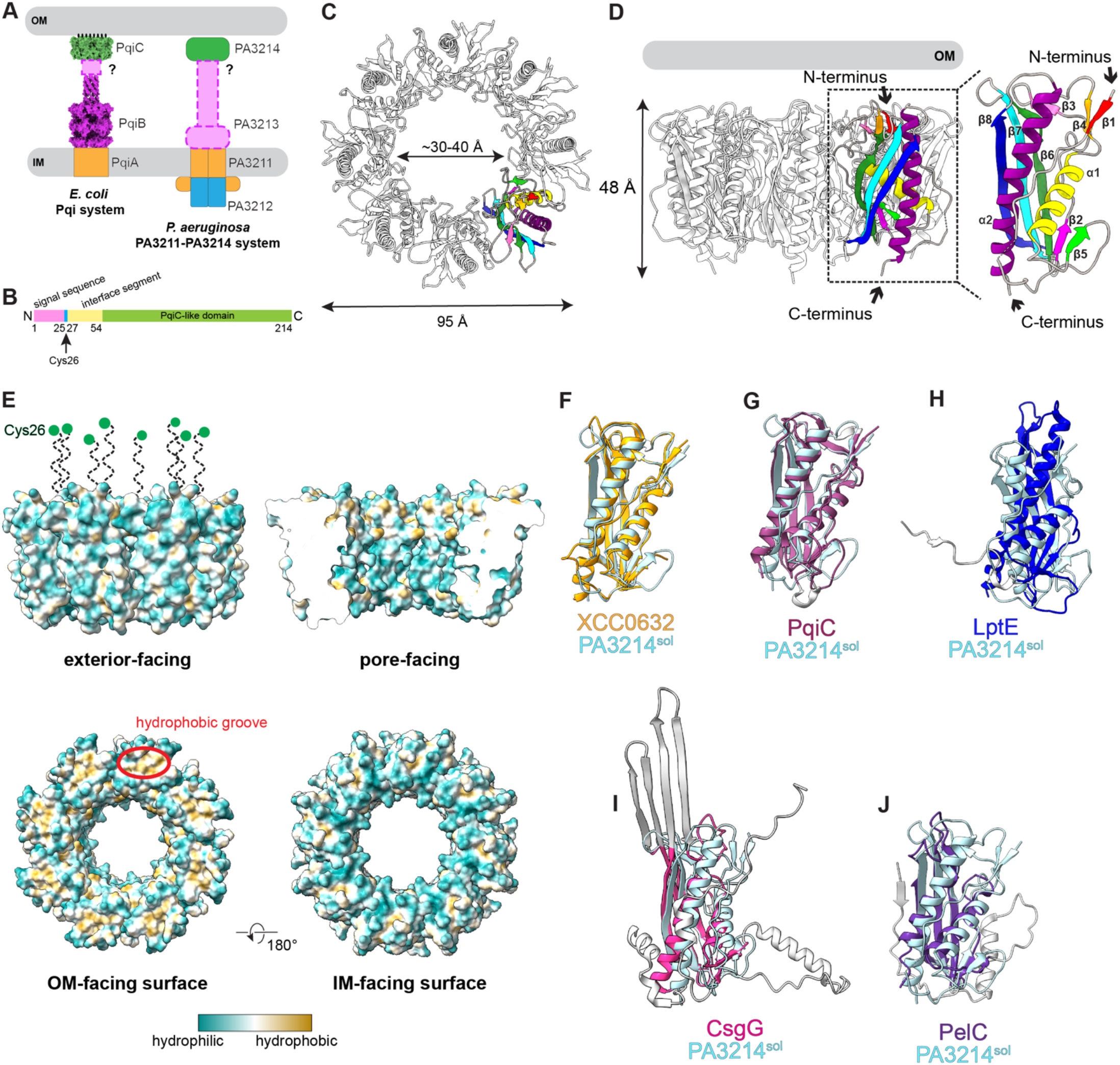
PA3214^sol^ lipoprotein forms an OM-anchored octamer with a central pore. *A*, model of the *E. coli* Pqi (PDB 5UVN, 8Q2C) MCE system and a schematic representation of the *P. aeruginosa* PA3211-PA3214 MCE system. These represent tripartite, trans-envelope systems with 1) an IM component (orange: transmembrane PqiA and PA3211; blue: ATPase PA3212), 2) an elongated, periplasmic MCE component (magenta: PqiB and PA3213), and 3) an OM lipoprotein component (green: PqiC and PA3214). IM and OM shown in gray, lipid anchors shown as black ovals. *B*, schematic representation of the PA3214 protein domain organization. *C*, cartoon representation of the OM-facing view of the PA3214^sol^ structure obtained from cryo-EM (Map 1), with one protomer colored in rainbow. *D*, cartoon representation of side-view of PA3214^sol^ structure (Map 1) in relation to the OM (shown in gray). Right, enlargement of a PA3214^sol^ monomer with sequential secondary structures labeled and colored in rainbow. *E*, molecular surface representation of PA3214^sol^ colored by hydrophobicity (ChimeraX). One hydrophobic groove is indicated by a red circle. The N-terminal lipid anchor (Cys26, green dot) and subsequent 7 residues of the mature lipoprotein were not resolved in our density and are shown as dashed lines drawn approximately to scale. *F*, overlay of PA3214^sol^ monomer (light blue) with *X. campestris* XCC0632 (PDB 2IQI) (orange). *G*, overlay of PA3214^sol^ monomer (light blue) with PqiC monomer (PDB 8Q2C) (dark magenta, with protein-specific regions in light gray). *H*, overlay of PA3214^sol^ monomer (light blue) with LptE (PDB 4N4R) (blue, with protein-specific regions in light gray). *I*, overlay of PA3214^sol^ monomer (light blue) with CsgG monomer (PDB 7BRM) (pink, with protein-specific regions in light gray). *J*, overlay of PA3214^sol^ monomer (light blue) with PelC monomer (PDB 9H80) (purple, with protein-specific regions in light gray).

The OM lipoprotein PqiC is encoded by the <u>P</u>ara<u>q</u>uat Inducible (Pqi) operon along with an IM transmembrane protein, PqiA (homolog of LetA (17)) and a periplasmic MCE protein, PqiB (13, 43) (**Fig. 1A**). PqiB resembles a needle-and-syringe, with three stacked MCE rings forming a barrel, and an elongated, ⍺-helical segment protruding from the barrel, creating the needle (13). This assembly is proposed to form a long, hollow tunnel, in which the interior of the tunnel is shielded from the aqueous periplasm, potentially allowing for translocation of lipids through the tunnel, across the periplasm (13). The structures of PqiC (42) and several PqiC-like proteins from other MCE systems (PDB codes: 2IQI and 6OSX) have been determined in isolation, in different oligomeric states. Full-length PqiC forms an octameric, ring-shaped assembly that is located predominantly in the periplasm and is predicted to associate with the OM via lipid anchors (42). A purified PqiAB complex has been shown biochemically to bind to PqiC, suggesting that PqiB and PqiC may interact (42), which is also supported by yeast 2-hybrid assays (22). A structure of the PqiBC complex is lacking, and the C-terminal region of PqiB that is hypothesized to interact with PqiC was not well resolved in the experimental structure (13), leaving open the question of how PqiB and other MCE proteins interact with PqiC-like OM lipoproteins (**Fig. 1A**).

In the human pathogen, *Pseudomonas aeruginosa*, the *PA3211-PA3214* operon encodes a PqiC-like OM lipoprotein, PA3214. While the other components of the PA3211-PA3214 system differ considerably from PqiABC, the C-terminal helical domains of the MCE proteins from both systems (PA3213 and PqiB) form homologous needle-like tubes, which may interact with their lipoprotein partners in similar ways. Consequently, we set out to characterize both the PqiABC and PA3211-PA3214 systems, to better understand how these MCE-lipoprotein pairs may interact. Here, we report the structures of both the soluble periplasmic portion of *P. aeruginosa* PA3214, as well as membrane anchored PA3214 bound to PA3213, highlighting interactions between these two proteins. Using a phenotype previously described for the homologous Pqi system (43), we use deep mutational scanning (DMS) to map the functionally important regions of PqiC. Structural comparisons of PqiC and PA3214, combined with DMS analysis, allow us to identify important sites of protein-protein interaction between the MCE protein and OM lipoprotein, which we test using biochemical assays in both systems. The key assembly interfaces between MCE proteins and their corresponding OM lipoproteins that we identify for the *E. coli* and *P. aeruginosa* systems likely represent principles that extend to other MCE systems that contain this unique conformation of elongated, needle MCE proteins and PqiC-like ring shaped OM lipoproteins.

## Results

### Lipoprotein PA3214 forms an homo-octameric ring

To resolve the structure and oligomeric state of PA3214, we expressed and purified a soluble PA3214 construct (PA3214^sol^), in which the first 26 residues were deleted (including the signal sequence and lipid attachment site, Cys26, **Fig 1B**), and determined its structure using single particle cryo-electron microscopy (cryo-EM). By size-exclusion chromatography, PA3214^sol^ elutes with an apparent molecular weight of ∼285 kDa, much larger than the expected molecular weight of a monomer (22.8 kDa), suggesting that PA3214^sol^ forms an oligomeric complex (**Fig. S2A**). Cryo-EM micrographs and 2D classes reveal particles consistent with ring-shaped oligomers (**Fig. S2B**). Most particles correspond to two PA3214^sol^ rings non-specifically interacting with each other (See Methods). We resolved the structure of a single PA3214^sol^ ring to an average resolution of 2.67 Å with C8 symmetry imposed (**Fig. 1C, Fig. S3, Fig. S4, Table S1**). Our final model of PA3214^sol^ is mostly complete, with only residues 27-34 and 46-51 unresolved in our EM map. We find that PA3214^sol^ is an octamer, with eight copies of PA3214^sol^ associating to form a ring with 8-fold rotational symmetry, surrounding a central channel (**Fig. 1C**).

PA3214 is predicted to be anchored to the OM via lipidation of the N-terminal Cys26 of the mature protein. The N-termini of all eight protomers lie on one face of the PA3214^sol^ ring, suggesting that the N-terminal face of the ring is oriented towards the OM (**Fig. 1D**). The C-terminus of each protomer is located at the opposite face of the ring, facing towards the periplasm and IM (**Fig. 1D**). The OM-facing surface contains a small, hydrophobic groove on each protomer (**Fig. 1E**), which may serve as a binding site for proteins or lipids. Barring the hydrophobic groove, the PA3214^sol^ surface is largely hydrophilic, suggesting that apart from the lipid anchors, PA3214^sol^ is unlikely to embed into the OM (**Fig. 1E**). Instead, we propose that the PA3214 ring is docked in close proximity to the OM, primarily held in place there via eight lipid anchors (coupled to Cys26 of each protomer). The PA3214^sol^ ring is 48 Å in height, with a ∼95 Å outer diameter and a pore diameter ranging from ∼30 - 40 Å (**Fig. S2C**). A single PA3214^sol^ protomer is composed of a mixed α/β fold with three β-strands (β6–β8) forming the core of a long, antiparallel β-sheet on one face of the protein, with a pair of α helices (α1 and α2) packing against the opposite side of the sheet (**Fig. 1D**). Two additional short, parallel strands (β2 and β5) extend the main sheet at one end. These two short strands and the coil following β5 project into the pore of the PA3214 ring near the IM-facing surface, forming a more constricted diameter at this section of the pore. IM-facing surface, forming a more constricted diameter at this section of the pore.

To explore structural similarity between PA3214^sol^ and previously determined structures, we performed a structure-based search of the PDB using FoldSeek (44) to identify any similarity to known protein folds (**Table 1**). As anticipated, the top hits are PqiC-like proteins (**Fig. S2D**): *X. campestris* XCC0632 (PDB 2IQI) (**Fig. 1F**); *E. cloacae* ECL_02694 (PDB 6OSX), which were deposited in the PDB by structural genomics centers without associated publication; and the *E. coli* lipoprotein PqiC (PDB 8Q2C) (42) (**Fig. 1G**). XCC0632 is the most structurally similar to PA3214 and is also an octamer, with a sequence identity of 33.5% (**Table 1**) (**Fig. S2E**,**F**). It is encoded in a four gene operon homologous to *PA3211-PA3214* (**Fig. S2G**). In contrast, the crystal structure of ECL_02694 is monomeric. This protein has a sequence identity of 13.5% to PA3214 and is encoded in an operon with *pqiA* and *pqiB*, homologous to the *pqiABC* operon in *E. coli* (**Fig. S2G**). *E. coli* PqiC is also more divergent, with a sequence identity of 12.6% and is encoded in an operon with substantially different MCE protein and IM membrane protein partners (**Fig. S2E, G**). Despite the sequence divergence, the structure of full-length PqiC forms an octameric ring assembly very similar to that of PA3214, and individual monomers are highly conserved structurally (42) (**Fig. 1G, Fig. S2H**). The OM-facing surface of both PqiC and XCC0632 also contain small, hydrophobic grooves similar to PA3214^sol^ (**Fig. 1E, Fig. S2F**). PqiC-like proteins are always associated with MCE transport systems and would be predicted to form octameric rings like PA3214, XCC0632 and PqiC.

**Table 1.**
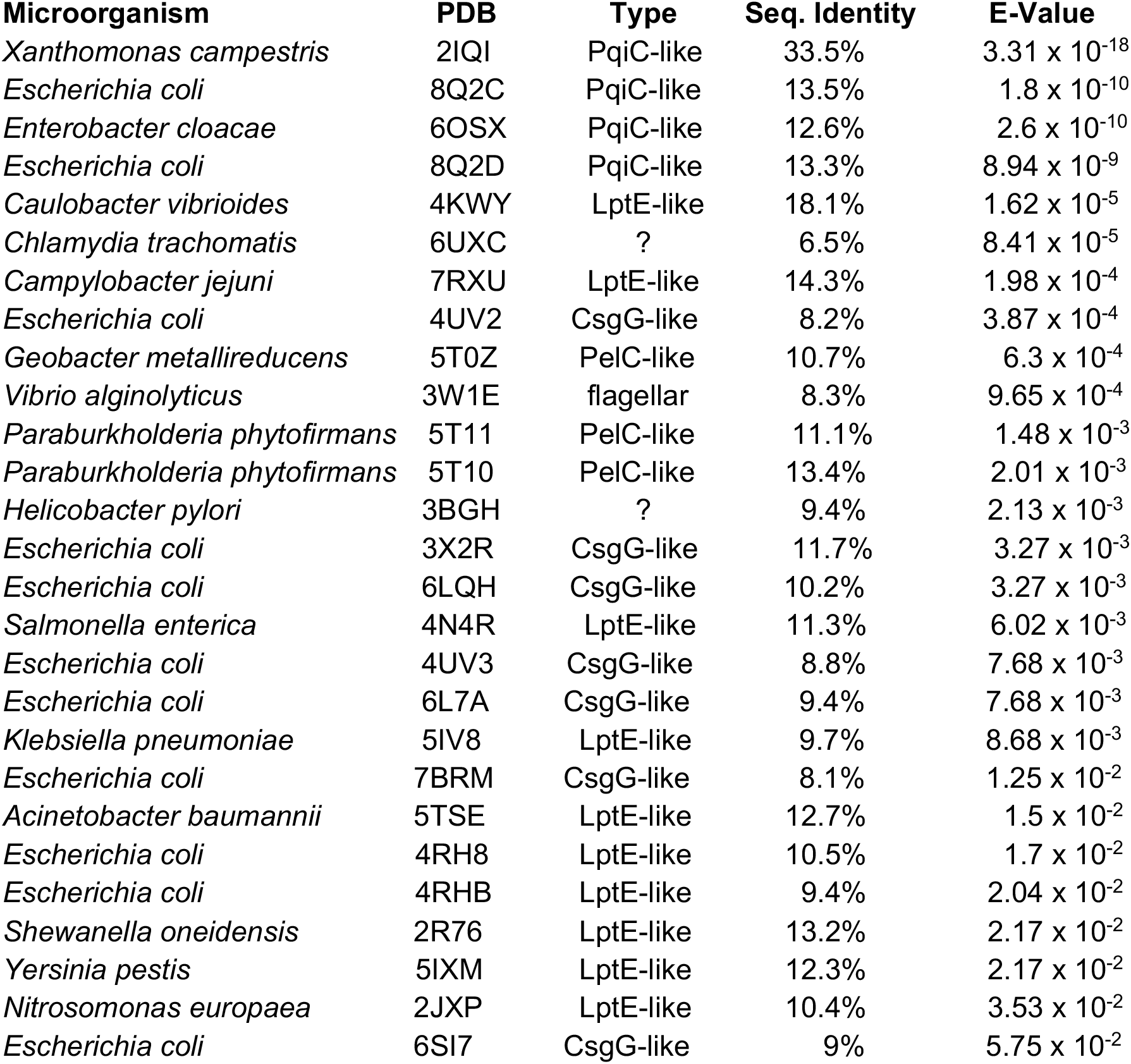
Structural homologs of *P. aeruginosa* PA3214.

Further hits from our Foldseek analysis could be categorized into three main groups: LptE-like, PelC-like, and CsgG-like proteins. All of these groups are OM lipoproteins that share a conserved α/β domain at their core (**Fig. S2I**), but differ in key aspects of their structures and the mechanisms by which they mediate transport across the bacterial cell envelope. First, in the LPS export system, a single copy of the LptE “plug” inserts into the β-barrel LptD (45–48) (**Fig. 1H, Fig. S2J**), in contrast to the octameric ring formed by PA3214^sol^. Thus, it is likely that PA3214 and LptE facilitate substrate translocation via distinct mechanisms. In contrast, CsgG and PelC are more similar to PA3214 in that they all form homooligomeric rings. *E. coli* CsgG forms a nonameric ring that mediates the secretion of the curli protein across the OM (49, 50) (**Fig. 1I, Fig. S2K**). The PqiC-like domains of CsgG associate with the periplasmic face of the OM, and each subunit contributes four β-strands to a 36-stranded β-barrel that creates a pore across the OM. However, the regions of CsgG that form the OM-spanning pore are absent in PqiC-like proteins (**Fig. 1I, Fig. S2I**,**K**), indicating that PA3214 would be unable to create an OM pore in a similar way. Finally, PelC, part of the Pel export pathway, forms a dodecameric ring involved in the translocation of Pel exopolysaccharide across the OM (51–53) (**Fig. 1J, Fig. S2L**). Like PA3214, the PelC ring resides in the periplasm, and lacks the membrane-spanning segments seen in CsgG. Intriguingly, the PelC ring interacts with an OM porin called PelB, potentially creating a pathway for substrate translocation through the center of the PelC ring and into the lumen of the porin and across the OM (54) (**Fig. S2L**). While PA3214/PqiC are not known to interact with porin proteins, the structural parallels with PelC raise the possibility that PA3214/PqiC could interact with porins in a similar way to facilitate lipid extraction or insertion at the OM, or perhaps even substrate translocation across the membrane.

### PA3213-PA3214 form a subcomplex long enough to span the periplasm

The octameric configuration of PA3214^sol^, XCC0632 (PDB 2IQI) and full-length PqiC (PDB 8Q2C) (43) is somewhat unexpected, as these proteins are all thought to interact with hexameric MCE proteins (13, 15, 16). The interaction between a hexameric MCE protein and octameric OM lipoprotein would result in a symmetry mismatch, and the structural basis of this interaction remains unknown. To determine the oligomeric state of PA3214 in the context of PA3213 and probe the PA3213-PA3214 interaction site, we pursued the biochemical and structural characterization of PA3214 following over-expression of the entire PA3211-PA3214 operon. We expressed the PA3211-PA3214 operon heterologously in *E. coli*, and performed affinity purification via a His-tag on the N-terminus of PA3211 (**Fig. 2A, Fig. S5A**). We found that the resulting sample elutes at an apparent molecular weight of ∼780 kDa by size exclusion chromatography, representing a much larger complex compared to PA3214^sol^ (∼285 kDa).

**Figure 2.**
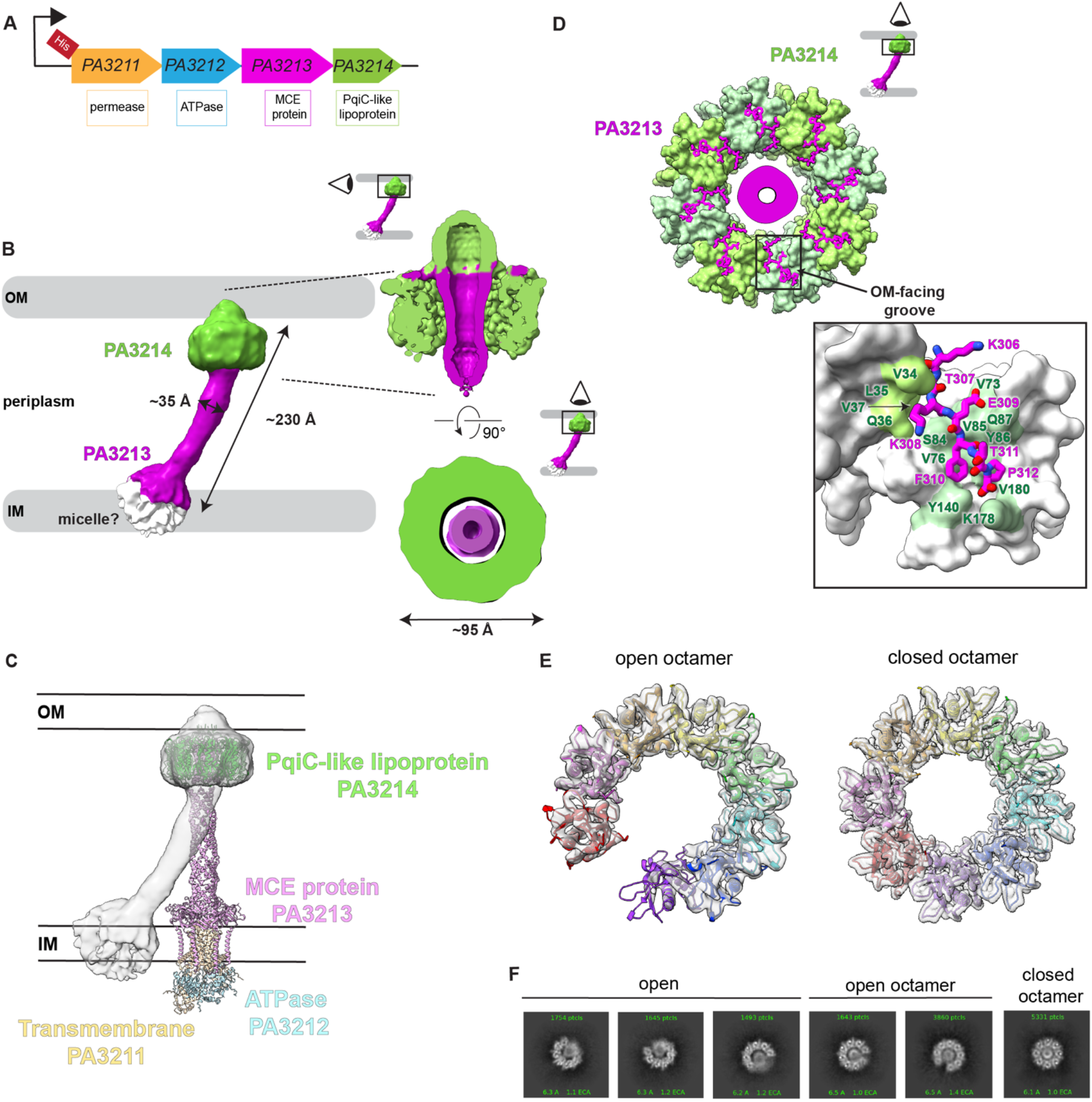
Structure of PA3214 OM lipoprotein bound to PA3213 MCE protein. *A*, schematic of the *P. aeruginosa PA3211-PA3214* operon overexpression construct used for structural studies. *B*, consensus map (Map 4) of protein complex after affinity purification and size exclusion. Inset shows a slice through the side view and top view of the consensus map, in which density likely corresponding to the PA3213 MCE protein is observed inside the PA3214 lipoprotein pore. *C*, AlphaFold 3 prediction of the PA3211-PA3212-PA3213-PA3214 complex, with PA3214 fit into the consensus map (Map 4). *D*, Surface representation of the PA3214 structure after focused refinement (Map 2, Fig. S6), with C-terminal peptide of PA3213 shown as sticks. Boxed region shows the interaction between the C-terminal peptide of PA3213 and the hydrophobic groove of PA3214. *E*, top views of the PA3214 model in the closed and open conformations fit into the respective maps (Map 2, closed; Map 3, open). Each protomer is shown in a different color. *F*, examples of 2D classes representing likely compositional and conformational heterogeneity in PA3214 bound to PA3213.

We characterized the resulting complex using single particle cryo-EM (**Fig. S5B, Fig. S6, Fig. S7A, Fig. S8, Table S2**), and obtained an initial low resolution reconstruction (7-16 Å). The map resembles a barbell, with two distinct globular masses ∼75-100 Å in diameter at either end of a ∼35 Å-thick rod that is ∼170 Å in length (**Fig. 2B**). To interpret our map, we predicted the structure of the complete PA3211-3214 complex using AlphaFold 3 (55) (**Fig. S5C**). Overall, AlphaFold 3 predicts a possible complex for PA3211-PA3214, with higher confidence in some regions, including the PA3211-PA3212 IM subcomplex, the PA3213 MCE ring, and the PA3214 octamer. The prediction of the ⍺-helical needle domain of the MCE protein, PA3213, has lower confidence and correspondingly, is substantially more variable (**Fig. S5C**). Compared with our EM reconstruction (**Fig. 2C**), several global elements of the PA3211-PA3214 AlphaFold model are in agreement, including the general shape of a long, helical needle from PA3213, reminiscent of the helical domains of Mce1 (16) and PqiB (13), with additional proteins bound at each end of the “barbell”. One end of the cryo-EM map is consistent with the presence of an octameric ring of PA3214 at the “OM end” of the needle, while the density at the other end, where we would predict the IM complex of PA3211-PA3212-PA3213 to be, is largely featureless (aside from the MCE ring domain). Further focused refinement of the putative IM-complex region did not reveal any clear protein features, raising the possibility that PA3211 and/or PA3212 has disassociated from the rest of the complex, and the density represents an empty detergent micelle (**Fig. 2B**). The long rod-like density connecting the two ends as well as the density for a ring of MCE domains are both consistent with a PA3213 hexamer as expected (**Fig. S5D**). The PA3213 α-helical needle is relatively straight in most AlphaFold predictions, but curved in our density map. The curvature of the PA3213 needle may represent a physiological conformation, or may represent the absence of the ABC transporter formed by PA3211-PA3212, which would anchor it in the IM (**Fig. S5C**). Overall, our map shows a periplasmic subcomplex ∼230 Å in length (measured from the bottom of the MCE ring to the top of PA3214), which is approximately consistent with the width of the periplasm (56–58).

### PA3213-PA3214 interaction interface

Focused refinement of the region corresponding to PA3214 resulted in a reconstruction of full-length PA3214 bound to PA3213 at an average resolution of 2.96 Åwith no symmetry applied (based on the expected symmetry mismatch between PA3213 and PA3214). While the regions of the map corresponding to PA3214 revealed considerable structural detail, the density for PA3213 lacked clear signs of secondary structure, possibly reflecting configurational heterogeneity of PA3213 relative to PA3214. To better resolve the features of PA3214 only, C8 symmetry was applied in the last round of refinement, resulting in an average resolution of 2.7 Å(residues Q33-P210 resolved in our map) (**Fig. 2D, Fig. S6, Fig. S7B, Table S2**). Full-length PA3214 remains an octamer when bound to PA3213 (**Fig. S5E**), and closely resembles PA3214^sol^ in isolation (root-mean square displacement (RMSD) of 0.472 Åacross C⍺ of 172 paired atoms). In contrast to the smaller and poorly defined densities in the PA3214^sol^ pore, a hollow tube of density runs through the center of the PA3214 ring in the full complex map (**Fig. 2B**), likely corresponding to the PA3213 needle. The map and our interpretation are consistent with AlphaFold predictions of the PA3214 ring assembling around the PA3213 needle (**Fig. 2B, 2D, Fig. S5C**). Despite considerable efforts to reconstruct the density inside the pore of PA3214 to higher resolution, we were unsuccessful, suggesting considerable flexibility between PA3213 and PA3214. One possibility is that the PA3213 needle adopts an ensemble of positions relative to PA3214, perhaps even freely rotating inside the PA3214 pore, like a spindle. Another possibility is that the PA3214 ring tilts, rotates and slides relative to the MCE needle, which is consistent with 3D classification and 3D variability analysis (See Methods, **Movies S1-3**). The different modes of flexibility are not mutually exclusive, and likely arise, to some extent, from the symmetry mismatch between PA3213 and PA3214 (6:8).

In both the C1 and C8 maps of PA3214 bound to PA3213, we observed additional density consistent with an extended, seven-residue peptide bound to the hydrophobic groove on the OM-facing surface of PA3214 (**Fig. 2D, Fig. S5F**). Based upon side-chain features, we assigned this density as the C-terminal seven residues of PA3213 (K306-P312) (**Fig. 2D**). This PA3213 C-terminal peptide nestles into the hydrophobic groove on the OM-facing surface of PA3214, which is formed by residues on two neighboring protomers (**Fig. 2D**). Notably, due to the symmetry mismatch between PA3213 and PA3214, there are eight potential binding sites on PA3214 but only six C-terminal PA3213 peptides. The presence of density for the C-terminal peptides at each of the eight binding sites in our C1 map with no symmetry applied suggests that each peptide may dynamically sample multiple binding sites, or that our density represents an average of multiple stable, distinct binding configurations, which we were unable to separate through our attempts at classification (see Methods).

Interestingly, 3D classification revealed two conformations of PA3214 octamer: a closed conformation in which the eight copies of PA3214 form a complete, closed ring and an open conformation in which eight copies of PA3214 are present, but the ring is not closed (**Fig. 2E, Fig. S6, Fig. S7C, Table S2**). Open oligomeric conformations with six visible copies of PA3214 were also observed in 2D classes during the pre-processing steps (**Fig. 2F**), suggesting that the PA3214 ring may assemble and/or disassemble around the PA3213 needle, as opposed to the needle threading through a pre-formed ring (**Fig. S5G, Fig. S7C**). Both the hexameric and octameric (open and closed) configurations of PA3214 are associated with the tip of the PA3213 needle. We hypothesize that the open conformation with six protomers is a compositional intermediate, and the open conformation with eight protomers is a conformational intermediate in the OM lipoprotein assembly or disassembly process. The final conformation is likely the closed, octameric PA3214 ring assembled around the PA3213 needle.

### Deep mutational scanning of PqiC identifies functionally important regions

The function of both *P. aeruginosa* PA3214 and its structural homolog *E. coli* PqiC remain poorly characterized. While PA3214 does not have a known phenotype, Δ*pqiC* strains display growth defects in the presence of the detergent lauryl sulfobetaine (LSB) (42, 43) (**Fig. 3A**). We took advantage of this phenotype to perform deep mutational scanning (DMS) on PqiC, to gain insights into functionally important sites in the PqiC-like family of OM lipoproteins. We constructed a plasmid library consisting of 5,984 mutants, in which each amino acid in PqiC (residues 1-187) was mutated to every other amino acid. This library was transformed into the Δ*pqiC* mutant strain and then grown in the presence of LSB, which enriches for functional PqiC variants. Using next-generation sequencing, we calculated the relative frequency of each *pqiC* mutant after LSB selection versus a non-selective medium and derived a comprehensive fitness landscape for PqiC based on the impact of each mutation (**Fig. S9A**). While many residues in PqiC tolerated mutations, 76 positions (∼40%) were more sensitive to LSB when mutated (based on a Tolerance Score [TS] < 0.9, where TS=0 indicates every possible mutation has an effect and TS=1 indicates mutations have no effect) (59–61) (**Fig. 3B, Fig. S9A, Table 2**). Of these 76 sensitive positions, seven residues are in the signal peptide which directs the secretion, translocation and anchoring of the lipoprotein to the OM (**Table 2**). 30 additional positions are located in the hydrophobic core of the protein (**Fig. S9B**), and are likely important for protein folding. The remaining 39 residues may represent functionally important sites in PqiC, and can be grouped into three spatially distinct clusters (**Fig. 3C**): (1) the oligomerization interface between PqiC protomers (22 residues); (2) the pore of the octameric ring (10 residues); and (3) the potential binding site for the C-terminal peptides from PqiB/PA3213 (7 residues).

**Table 2.** PqiC DMS Scores.

| **Protomer-protomer interface** | <u>Position</u> | <u>Residue</u> | <u>Tolerance Score</u> |
| --- | --- | --- | --- |
|  | 26 | Y | 0.53846154 |
|  | 27 | Q | 0.48611111 |
|  | 30 | V | 0.77380952 |
|  | 54 | P | 0.64893617 |
|  | 56 | Y | 0.73076923 |
|  | 65 | Q | 0.625 |
|  | 67 | S | 0.897727273 |
|  | 69 | V | 0.53571429 |
|  | 70 | K | 0.86666667 |
|  | 80 | A | 0.21428571 |
|  | 81 | S | 0.20454545 |
|  | 92 | V | 0.80952381 |
|  | 96 | S | 0.86363636 |
|  | 103 | V | 0.53571429 |
|  | 105 | A | 0.70238095 |
|  | 106 | S | 0.89772727 |
|  | 121 | T | 0.72058824 |
|  | 122 | E | 0.82653061 |
|  | 134 | S | 0.27272727 |
|  | 138 | L | 0.40243902 |
|  | 145 | L | 0.84146341 |
|  | 147 | K | 0.4 |
| **Pore-facing** | 23 | K | 0.78888889 |
|  | 55 | D | 0.64 |
|  | 58 | A | 0.72619048 |
|  | 59 | G | 0.67857143 |
|  | 60 | N | 0.7972973 |
|  | 61 | G | 0.4047619 |
|  | 73 | I | 0.75609756 |
|  | 78 | L | 0.890243902 |
|  | 82 | P | 0.60638298 |
|  | 85 | Q | 0.56944444 |
| **OM-surface facing** | 64 | Y | 0.73076923 |
|  | 74 | A | 0.47619048 |
|  | 77 | N | 0.7972973 |
|  | 129 | G | 0.45238095 |
|  | 160 | G | 0.70238095 |
|  | 161 | Y | 0.35897436 |
|  | 164 | M | 0.825 |
| **Signal peptide** | 1 | M | 0.325 |
|  | 8 | I | 0.76829268 |
|  | 10 | A | 0.8452381 |
|  | 11 | L | 0.70731707 |
|  | 13 | L | 0.86585366 |
|  | 15 | G | 0.75 |
|  | 16 | C | 0.51470588 |
| **Hydrophobic core** | 28 | L | 0.2804878 |
|  | 46 | L | 0.58536585 |
|  | 48 | V | 0.58333333 |
|  | 51 | V | 0.64285714 |
|  | 53 | V | 0.67857143 |
|  | 57 | L | 0.17073171 |
|  | 62 | V | 0.60714286 |
|  | 63 | V | 0.71428571 |
|  | 79 | W | 0.32051282 |
|  | 83 | L | 0.43902439 |
|  | 86 | Q | 0.84722222 |
|  | 87 | L | 0.37804878 |
|  | 91 | L | 0.67073171 |
|  | 95 | L | 0.59756098 |
|  | 104 | V | 0.84615385 |
|  | 116 | L | 0.5 |
|  | 118 | V | 0.63095238 |
|  | 120 | V | 0.60714286 |
|  | 123 | F | 0.525 |
|  | 124 | N | 0.85135135 |
|  | 125 | G | 0.55952381 |
|  | 131 | V | 0.72619048 |
|  | 133 | V | 0.66666667 |
|  | 135 | G | 0.1547619 |
|  | 150 | F | 0.6875 |
|  | 165 | V | 0.47619048 |
|  | 168 | L | 0.7195122 |
|  | 172 | W | 0.30769231 |
|  | 176 | A | 0.8452381 |
|  | 179 | I | 0.7195122 |

**Figure 3.**
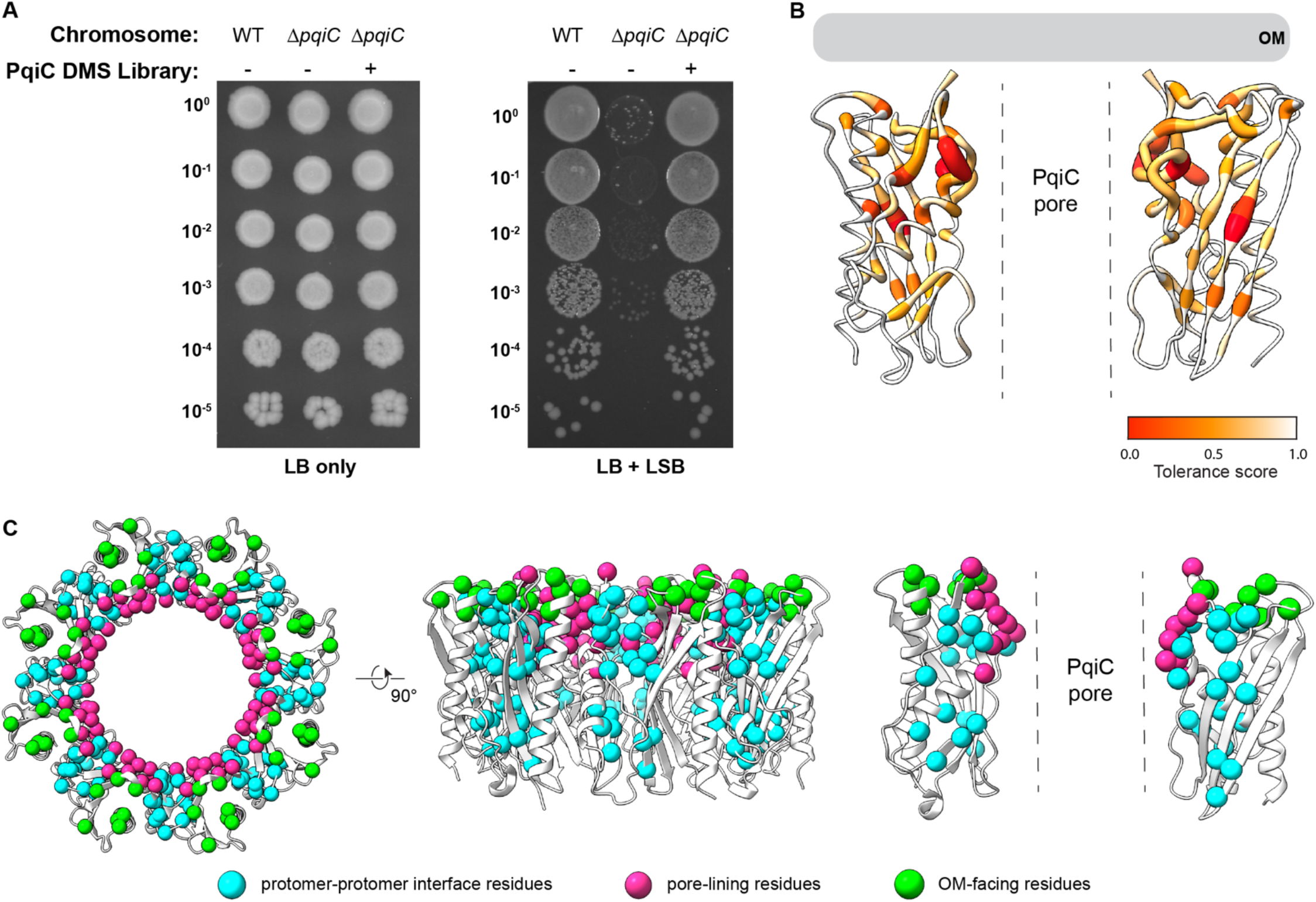
Deep mutational scanning points to functionally important regions of PqiC. *A*, cellular assay to assess the function of the PqiC DMS library. Ten-fold serial dilutions of strains were spotted on LB agar with or without LSB. *B*, protomers of PqiC colored by tolerance scores from DMS analysis. Residues most sensitive to mutation are the deepest shade of red and are represented by the thickest backbone trace. *C*, cartoon representation of octameric PqiC, where residues with tolerance scores <0.9 (excluding signal peptide, lipid anchor, and protein core) are depicted by spheres at each C⍺ position. Three spatially distinct regions are highlighted: protomer-protomer interface (cyan), pore-facing residues (pink), and OM-facing residues (green).

#### Protomer-protomer interface

22 mutation-sensitive residues lie at the interface between adjacent PqiC protomers, suggesting a role in protomer-protomer interaction and oligomerization of the PqiC ring (**Fig. 4A,B, Table 2**). Of the 22 residues that cluster to the interface between PqiC protomers, ten are located on one face of the protomer, consisting of helix α1, strands β1 and β5, and the “interface segment” (a coil immediately following the N-terminal lipid anchor and linker, which is sandwiched between adjacent lipoprotein protomers). The remaining residues are located on the other face of the protomer, at the OM proximal extended loop and the beta sheet (β6-β8) (**Fig. 4A**). Three of the mutation-sensitive interface residues are specifically located on the interface segment (residues Tyr26, Gln27, Val30) (**Fig. 4A**). To test whether this interface segment plays a role in oligomerization, we deleted the interface segment from our PA3214^sol^ construct, yielding PA3214^Δintseg^ (**Fig. 4C**), and used size exclusion chromatography to assess the oligomeric state. The size exclusion profile for PA3214^Δintseg^ is shifted towards much smaller molecular weights, with the main peak eluting at ∼17 kDa, corresponding to monomers and possibly small oligomers (**Fig. S9C**), compared to ∼285 kDa for PA3214^sol^. Consistent with this, negative stain EM shows larger particles for the construct that retains the interface segment (lPA3214^sol^), while these were not observed for PA3214^Δintseg^ (**Fig. S9D**). These results suggest that the N-terminal interface segment is involved in facilitating interactions between PqiC-like protomers.

**Figure 4.**
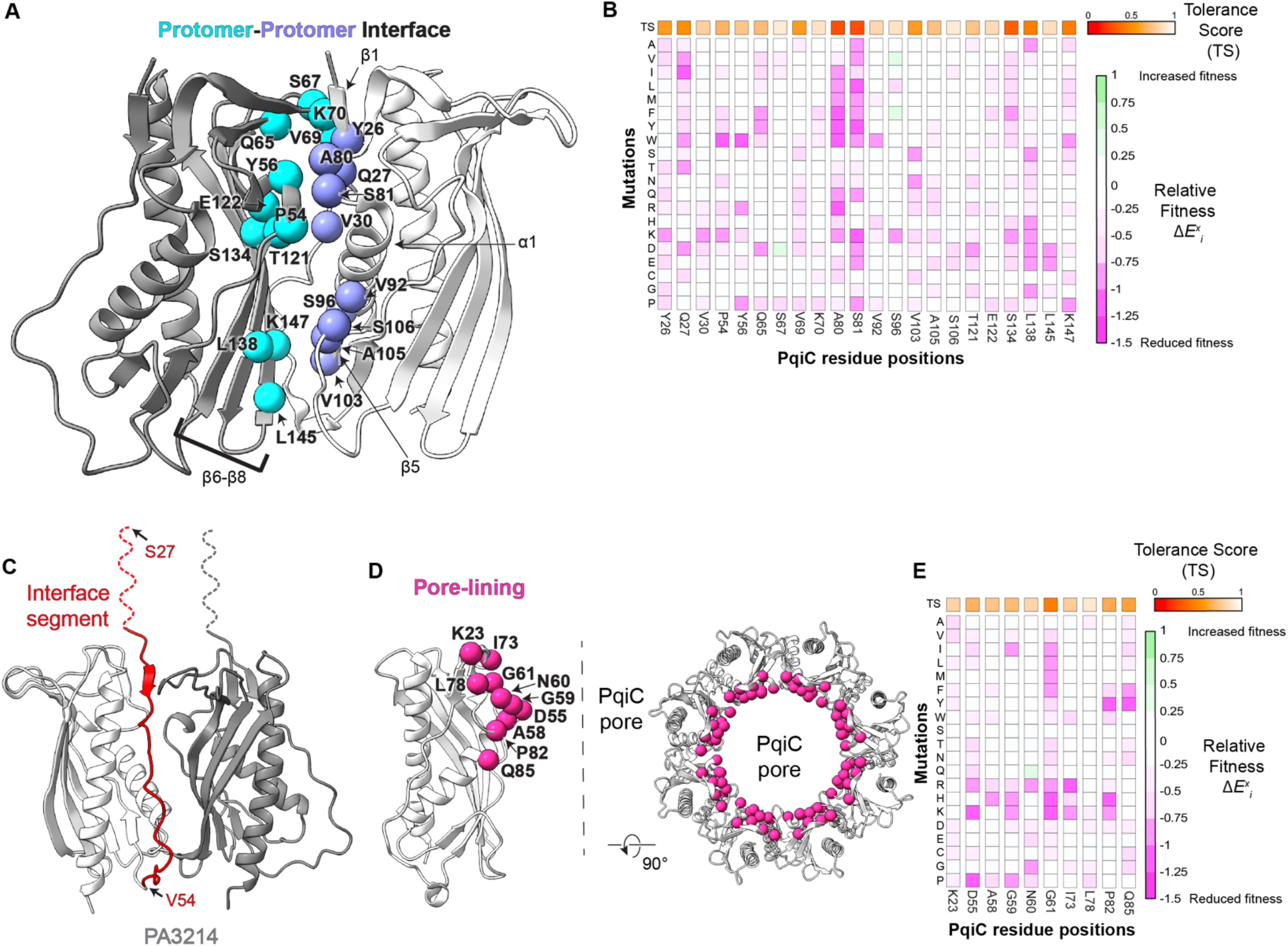
PqiC residues sensitive to mutation at the protomer-protomer interface and lining the pore. *A*, two neighboring PqiC protomers (white and gray) shown in cartoon representation, where interface residues identified as important by DMS are depicted by spheres at each C⍺ position. *B*, DMS data corresponding to the residues shown in *A*. Vertical strips corresponding to individual PqiC residues are from the heat map shown in Fig. S9A. Each square represents the average fitness cost of an individual mutation relative to the WT sequence from two biological replicates. Mutations that result in reduced fitness relative to the WT are shown in shades of magenta, while mutations that result in increased fitness are shown in shades of green. The Tolerance Score (see Methods) is shown as the colored square (red-to-white) above each strip. *C*, cartoon representation of two PA3214 protomers (white and gray), with interface segment in red. The portion of the interface segment for which density is not observed is shown as a red dotted line, drawn to scale. *D*, PqiC monomer (side view, left) and octamer (top view, right). Pore-lining residues identified as important by DMS are depicted by magenta spheres at each C⍺ position. *E*, DMS data corresponding to the residues shown in *D*, as presented in panel *B*.

#### Pore-facing

Of the ten mutation-sensitive residues that map to the PqiC pore, five of these residues lie in a loop that protrudes into the central pore (Asp55, Ala58, Gly59, Asn60 and Gly61; (**Fig. 4D,E, Table 2**). Based upon our AlphaFold prediction of the PqiBC interface (**Fig. S9E, S9F**), this loop may be interacting with residues (R513, E514, K520) on the helical needle of PqiB as it passes through the pore of PqiC. Alternatively, mutations in this loop could potentially influence the OM-facing peptide binding groove located immediately above. Pro82 and Gln85 are part of a pore-proximal loop as well, that extends into the central pore of PqiC to potentially interact with some residues on the PqiB needle (**Fig. S9F**). The last three mutation-sensitive residues that line pore of PqiC, Lys23, Ile73, and Leu78, are near the OM-facing surface of the lipoprotein. Ile73 and Leu78 protrude into the pore, placing these residues in proximity to the helical domain of PqiB needle and potentially facilitating interactions (**Fig. S9F**). Lys23 may interact with the very C-terminus of the helical domain of the MCE needle. Besides these ten mutation-sensitive residues, the rest of the positions that line the central pore of PqiC appear to be tolerant of most mutations, consistent with previous results (42). Consistent with the PA3211-PA3214 complex prediction, modeling of PqiBC suggests that the width of the PqiB MCE needle is ∼32 Å, leaving space between the needle and the wall of the PqiC pore (diameter ∼45 Å) that may minimize direct protein-protein contacts, and may allow for rotation of the MCE needle (**Fig. S9G**).

#### Binding site for MCE C-terminal peptides

Similar to the binding interactions observed in our experimental PA3213-PA3214 structure, our AlphaFold prediction of the PqiB:PqiC complex suggest that the helical needle of PqiB threads through the PqiC pore, and that the PqiB C-terminal peptides dock onto the OM-facing surface of the PqiC ring. This predicted interaction accounts for the final cluster of functionally important residues identified in our DMS experiment: Tyr64, Ala74, Asn77, Gly129, Gly160, Tyr161, and Met164 (**Fig. 5A-C, Table 2**). To better understand the protein-protein interactions between PA3213 and PA3214 as well as PqiB and PqiC, we designed mutations in both systems based on the DMS results of PqiC, and carried out pull-down experiments to assess their impact on binding between the two proteins. When His-tagged PA3214 was pulled down from *E. coli* cells over-expressing the entire *PA3211-PA3214* operon, PA3213 co-purified as part of the resulting complex. Similarly, when His-tagged PqiB was pulled down from *E. coli* cells over-expressing the entire *pqiABC* operon, PqiC co-purified. Deletion of the C-terminal peptide of each MCE protein (residues 296-312 in PA3213 and residues 530-546 in PqiB) has minimal impact on protein expression but disrupts binding in both systems, consistent with a role in mediating MCE-lipoprotein interactions (**Fig. 5D, E**). To probe the interactions further, we designed single amino acid substitutions in the PA3213 C-terminal peptide (E309G, F310G, P312G), which greatly diminish the interaction of PA3213 with PA3214 (**Fig. 5D**). Four out of five point mutations in the peptide binding site on PA3214 (R73E, V76K, Y86K, and V180K) diminish the interaction with PA3213 (**Fig. 5D**), while a K178E has little effect. Similar results were observed when mutations were introduced at the corresponding locations in PqiB and PqiC (PqiB mutations D538G, P539G, P541G and PqiC mutations Y64K and Y161K) (**Fig. 5E**). These data suggest that the interface between the MCE C-terminal peptide (PA3213/PqiB) and the hydrophobic groove on the OM lipoprotein (PA3214/PqiC) is critical for the interaction between these proteins. In contrast, the interface between the MCE needle and the pore of the lipoprotein ring seems to be relatively insensitive to mutation.

**Figure 5.**
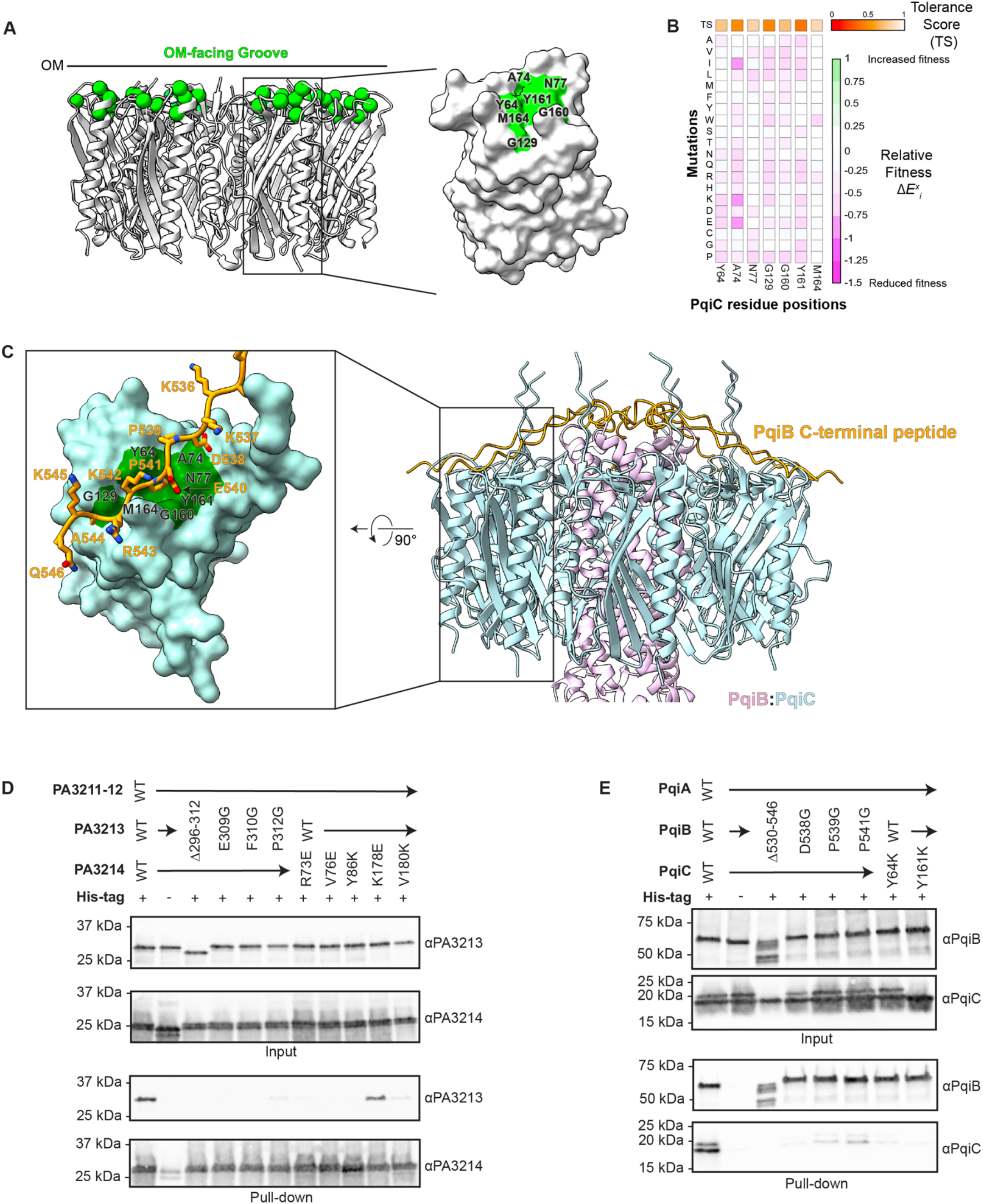
Interactions between MCE protein and OM lipoprotein. *A*, cartoon representation PqiC octamer and surface representation of a monomer, where hydrophobic groove residues identified as important by DMS are depicted in green on both the octamer and the monomer. *B*, DMS data corresponding to the residues shown in *A*, as presented in Fig. 4B. *C*, Alphafold 3 prediction of PqiC in complex with the C-terminal region of PqiB. Surface representation of the PqiC predicted model with C-terminal peptide of PqiB shown as sticks. Inset focuses on the predicted interaction between PqiB and PqiC, showing the C-terminal peptide of PqiB and surrounding regions in the hydrophobic groove of PqiC. *D*, representative Western blot from a pull-down assay to assess the interaction between PA3213 and PA3214. All four subunits of the PA3211-PA3214 complex were over-expressed, with a His tag on the PA3214 bait, and interaction with untagged PA3213 was assessed using an ⍺-PA3213 antibody. Blots showing the solubilized membrane fraction of each strain (input, expression control), and the results of the pull-down are shown. Three independent purifications were performed starting with three different inoculations, with similar results. *E*, representative Western blot from a pull-down assay to assess the interaction between PqiB and PqiC. All three subunits of the PqiABC complex were over-expressed, with a His tag on the PqiB bait, and interaction with untagged PqiC was assessed using an ⍺-PqiC antibody. Blots showing the solubilized membrane fraction of each strain (input, expression control), and the results of the pull-down are shown. Three independent purifications were performed starting with three different inoculations, with similar results.

## Discussion

Our data provide key insights into the interaction between MCE proteins and PqiC-like OM lipoprotein binding partners. The cryo-EM structure of *P. aeruginosa* PA3214^sol^, a homo-octameric ring (**Fig. 1C**), is in agreement with previously determined structures of homologs in isolation, including *E. coli* PqiC (42). The cryo-EM structure of full length PA3214 in the context of MCE protein PA3213 provides new insight into the interaction between these two proteins. While our highest resolution structure is of the PA3214 octamer, we also observe other conformations of PA3214 in this dataset. This includes an open conformation, in which the octameric ring is not fully closed, and possible compositional intermediates that appear to be open (**Fig. 2E,F**).

Using an unbiased DMS screen, we uncovered functionally important regions of the PA3214 homolog, PqiC (**Fig. 3B,C**), which supports the importance of a key site of interaction observed in our PA3213-PA3214 structure and PqiB-PqiC AlphaFold prediction. The C-terminal peptides of PA3213/PqiB interact with a hydrophobic groove on the OM-facing surface of PA3214/PqiC, providing a critical interface required for MCE protein binding to its cognate PqiC-like lipoprotein (**Fig. 6**). Mutations in key interface residues abolish binding between PA3213 and PA3214 or PqiB and PqiC. What remains less clear from currently available data is to what extent the α-helical needle of PA3213 or PqiB interacts with the lumen of the PA3214/PqiC ring. Our cryo-EM data remain at low resolution in this region: adequate resolution to confirm that the PA3213 needle lies within the lumen of the PA3214 pore, but too low to determine the interaction interface. The needle likely has a high degree of structural variability within the PA3214 pore, and better understanding the nature of this flexibility, specific interactions that may be formed, and how flexibility of transient interactions relate to function remains an open question.

**Figure 6.**
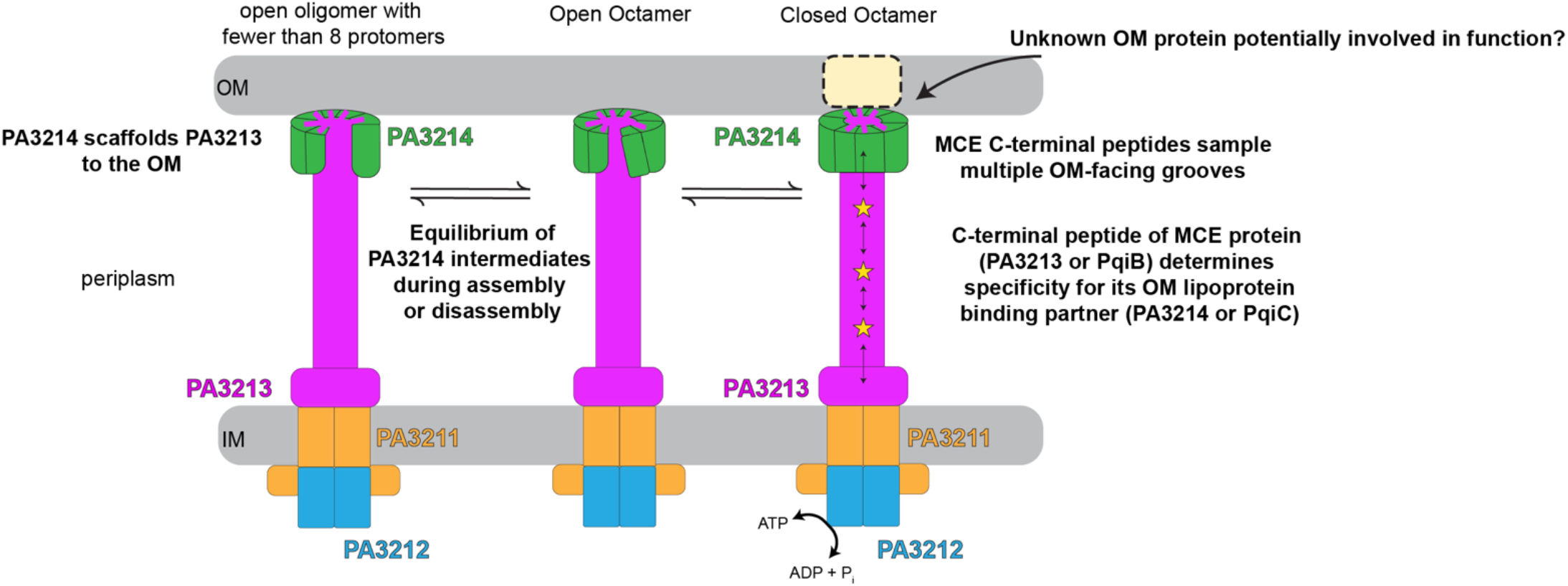
A model highlighting MCE protein-OM lipoprotein interactions in the context of the PA3211-PA3214 transport system. PA3214, and likely PqiC are compositionally and conformationally heterogeneous. Closed octamers that fully enclose PA3213 (PqiB) may represent the functional transport system, while open octamers and oligomers with fewer than 8 subunits may represent assembly or disassembly intermediates. The MCE protein needle, formed by the C-terminus of PA3213 or PqiB, may rotate freely within the PA3213/PqiC pore. The specificity of the interaction between the MCE protein and the OM lipoprotein is determined by the sequence of the MCE protein C-terminal tail, and the corresponding hydrophobic groove on the OM lipoprotein.

The proposed function of the PqiABC and PA3211-PA3214 MCE systems is to transport lipids, possibly phospholipids across the cell envelope. Our work provides insight into the interaction between the OM lipoprotein and MCE protein, but it remains unclear whether these systems transport substrates between the IM and OM, as is thought to be the case for the *E. coli* MCE system, Mla (62–65), or whether they transport substrates across the OM, analogous to the LPS (45, 47, 48, 63), Csg (49, 50) or Pel (51– 53) transport systems. In the Mla system, a monomeric, OM lipoprotein, MlaA, is scaffolded onto the OM embedded porin OmpF/C, and a channel formed within MlaA allows extraction of phospholipids from the OM (34). On the other hand, *P. aeruginosa* PelC, an OM lipoprotein, interacts closely with OM porin PelB, to form a continuous pathway for substrate translocation across the OM. In the case of MCE systems PA3211-3214, PqiABC and other homologs, it is still unclear how translocation through the OM may occur, if at all. One possibility is that the OM lipoproteins from these systems interact with unknown OM binding partners (**Fig. 6**), while another possibility is that PA3214, PqiC and other homologs directly deform the OM in ways that are yet to be defined, and may require near-native membrane reconstitution to address. Our structural and functional data provide a framework for understanding the interactions between MCE proteins and their OM PqiC-like lipoprotein binding partners, and the complex, dynamic nature of these interactions.

## Methods

### Expression and purification of PA3214^**sol**^

Plasmid pBEL2277, which encodes for N-terminal 6xHis2xQH-TEV tagged PA3214^sol^ (PA3214 residues 27-214) was transformed into Rosetta 2(DE3) cells (Novagen #714003) (Table S3). For protein expression, an overnight culture (LB + 100 µg/mL carbenicillin + 38 µg/mL chloramphenicol) was diluted 1:40 in LB (Difco) supplemented with carbenicillin (100 µg/mL) and chloramphenicol (38 µg/mL), grown at grown at 37°C with shaking to an OD600 of 0.6-0.8, and induced by the addition of L-arabinose (final concentration 0.2%). Cultures were then incubated at 37°C with shaking for 4 hours, and then harvested by centrifugation at 4,500g for 15 minutes. Pellets were resuspended in 50 mM Tris pH 8.0, 300 mM NaCl, and 1x cOmplete Protease Inhibitor (Roche). Cells were lysed by two passes through an Emulsiflex-C3 cell disruptor (Avestin), then centrifuged at 25,000g for 20 min at 4°C to pellet cell debris. The supernatant was incubated with Ni Sepharose resin (Cytiva 17-5318-02) for 30 min at 4°C on a rocker, and then washed with 50 column volumes of 50 mM Tris-HCl pH 8.0, 300 mM NaCl, 10 mM imidazole; then 100 column volumes of 50 mM Tris-HCl pH 8.0, 300 mM NaCl, 20 mM imidazole; and then 100 column volumes of 50 mM Tris-HCl pH 8.0, 300 mM NaCl, 40 mM imidazole to remove non-specifically bound proteins. The target protein was eluted with 20 mL 50 mM Tris pH 8.0, 300 mM NaCl, 250 mM imidazole in four fractions of 5 mL volume each. Eluted fractions containing the target protein were pooled and concentrated to 0.5 mL using an Amicon Ultra-15 Centrifugal Filter Unit concentrator (MWCO 10 kDa, UFC901024) before separation on the Superdex 200 Increase 10/300 column (Cytiva) equilibrated with HEPES (20mM HEPES pH 7, 150mM NaCl) size exclusion buffer.

### PA3214^sol^ cryo-EM sample preparation and data collection

Fractions from size exclusion containing PA3214^sol^ were pooled and concentrated to 0.3 mg/mL using an Amicon Ultra-0.5 centrifugal filter unit concentrator (MWCO 10 kDa, UFC501096). Quantifoil R1.2/1.3 on Cu 400 mesh grids were glow-discharged in a PELCO easiGlow Glow Discharge Cleaning System (Ted Pella) and 3.5 μL of freshly prepared sample was added to the glow-discharged grid. Grids were prepared using a Vitrobot Mark IV (Thermo Fisher Scientific): grids were blotted with a blot force of 0 for 6 s at 4°C with 100% chamber humidity and then plunge-frozen into liquid ethane. Grids were clipped for data acquisition and imaged at the Pacific Northwest Center for Cryo-EM (PNCC) on Krios-1, a Titan Krios G3 electron microscope (Thermo Fisher Scientific) equipped with a K3 direct electron detector with a BioContinuum energy filter (Gatan). Super-resolution movies were collected at 300 kV using SerialEM (66) at a nominal magnification of 22,500x, corresponding to a super-resolution pixel size of 0.5144 Å (or a nominal pixel size of 1.029 Å after binning by 2). 6,463 movies were collected over a defocus range of −0.8 to −2.1 µm and each movie consisted of 50 frames with a total dose of 50 e-/Å2. Data collection parameters are shown in **Table S1**.

### Cryo-EM data processing of the PA3214^sol^

The workflow associated with the processing of the soluble domain is described in Fig. S3. Movies were first aligned with Relion (67), using the in-built Relion implementation of motion correction, then transferred to CryoSPARC (68) for curation, and the 4,157 micrographs left were CTF-corrected using CTFFIND4 (69). Note that the large number of micrographs removed was not due to a sample quality issue, but the column valve was closed during the acquisition leading to hundreds of empty micrographs.

Particles were picked in CryoSPARC iterating through several rounds of picking/2D classification strategies. In detail: 37 micrographs were first randomly selected and 43,236 particles were picked searching for blobs from 100 to 200 Å. After extracting the particles with a box size of 220 pixels, three rounds of 2D classification followed by template picking were performed on various subsets of the 4,157 micrographs. The 47,638 particles from that last round were then fed into Topaz (70) where 3,329,863 particles were picked from all 4,157 micrographs and submitted to three rounds of 2D classification. Six templates from 346,047 particles were selected for a final round of template picking on the full dataset, resulting in 3,638,701 particles extracted with a box size of 150 pixels. After 2D classification, 1,882,105 were selected, while a subset of 142,235 junk particles were used to generate decoy models via the ab-initio job. During this process, initial 2D classes reveal ring-shaped oligomers, but with two PA3214^sol^ rings interacting with each other (**Fig. S3A**). We believe this to be due to non-specific interactions between the OM-facing surfaces and the his-tags of both PA3214^sol^ rings. In the initial classes, fuzzy 2D classes aligned on the two rings can be seen. To reach higher resolution for a single ring, an extensive iterative 2D classification strategy was used, tightening the box size from 220 pixels to 150 pixels. In the later classes, 2D classes show higher resolution features with one of the rings dominating the alignment and better centered in the 2D classes.

An initial ab-initio reference model was also obtained using a subset of particles from the second round of particle picking (**Fig. S3B**). We selected the 4 classes where both the size and the top view of the rings were properly centered and where the alignment was dominated by one of the rings. Template picking led to 7,017,815 particles extracted with a very tight box size (128 pixels), and subject to two rounds of 2D classification. One of the four ab-initio models obtained from the resulting 2,831,031 particles show a single well-centered ring. That ab-initio model was refined twice with homogeneous refinement, with C8 symmetry applied in the second round as well as a mask on the PA3214^sol^ ring.

The resulting model and two decoy models created from the 142,235 junk particles were then used as input for a 3-class heterogeneous refinement of the selected 1,882,105 total particles (**Fig. S3C**). The particles from the best resulting map were then subject to homogeneous refinement (with C8 symmetry), and the cycle between heterogeneous and homogeneous refinements was iterated 4 more times, until the resolution of the final homogeneous model stopped getting better. The final round of heterogenous refinement was performed with 5 classes, and 706,909 particles (about 95%) were selected, which led to a map with an average resolution of 2.81 Å after homogenous refinement.

The dataset was then transferred to Relion (**Fig. S3D**) via the pyem tools (71). The particles were first subject to a round of 2D classification followed by removal of duplicate particles. The 654,956 particles left were auto-refined to 2.97 Å before running the CTFRefine and the particle polishing tools, improving the Relion-estimated resolution to 2.81 Å. The data was then moved back to CryoSPARC for a final round of 3D non-uniform refinement with C8 symmetry and a static mask covering the ring density, leading to a final 2.67 Å estimated resolution (Map 1, **Table S1, Fig. S4**).

### Model building of PA3214^sol^

We first generated a homology model using Phyre2 (72) for the PA3214sol monomeric construct, as this structure was modeled prior to AlphaFold availability. Eight copies of the monomer were manually docked into the cryo-EM map using the fit_in_map tool in Chimera (73). Additional diffuse density was observed within the central pore, which we did not explicitly model, but which may derive from the His-tag of each PA3214^sol^ protomer from both rings. This octameric model was then used as the starting model for real-space refinement in Phenix 1.16 (74). The first round of refinement was done with the rigid-body option on, without restraints, and with each chain defined as an independent rigid body. Subsequent rounds were performed with Ramachandran, rotamer, NCS, and secondary structure restraints, with manual model adjustment in Coot (75) between each round of automated refinement. In later rounds of automated refinement, secondary structure restraints were turned off and ADP and reference model restraints were turned on. Because of the lack of density, we did not model residues 27-34 and residues 201-214, as well as residues 48-50 associated with the most flexible part of one of the loops.

### Expression and purification of PA3211-PA3214

Plasmid pBEL3076, which encodes N-terminally 6xHis2xQH-TEV-tagged PA3211 and untagged PA3212-PA3213-PA3214, was transformed in Rosetta 2(DE3) cells (Novagen #714003) (**Fig. 2A, Table S3**). For protein expression, an overnight culture (LB + 100 µg/mL carbenicillin + 38 µg/mL chloramphenicol) was diluted 1:50 in LB (Difco) supplemented with carbenicillin (100 µg/mL) and chloramphenicol (38 µg/mL), grown at grown at 37°C with shaking to an OD600 of 0.6-0.8, and induced by the addition of arabinose (final concentration 0.2%). Cultures were then incubated at 37°C with shaking for 4 hours, followed by harvesting by centrifugation at 4,500g for 20 minutes. Pellets were resuspended in 50 mM Tris pH 8.0, 300 mM NaCl, 10% glycerol, flash frozen in liquid nitrogen, and stored at −80°C. Cells were lysed by three passes through an Emulsiflex-C3 cell disruptor (Avestin), then centrifuged at 32,000g for 35 minutes at 4°C to pellet cell debris. The clarified lysate underwent ultracentrifugation at 37,000 rpm (182,460g) for 49 minutes at 4°C in a Fiberlite F37L-8 x 100 Fixed-Angle Rotor (Thermo Fisher Scientific, 096-087056). The supernatant was discarded and the membrane fraction was solubilized in 50 mM Tris pH 8.0, 300 mM NaCl, 10% glycerol, 25 mM n-DodecylB-D-maltoside (DDM) by rocking overnight at 4°C. Insoluble debris was pelleted by ultracentrifugation at 37,000 rpm for 49 minutes at 4°C. Solubilized membranes were incubated with Ni Sepharose resin (Cytiva 17-5318-02) for 30 min at 4°C, and then washed with 10 column volumes of 50 mM Tris-HCl pH 8.0, 300 mM NaCl, 10 mM imidazole, 0.5 mM DDM and 10% glycerol; then 20 column volumes of 50 mM Tris-HCl pH 8.0, 300 mM NaCl, 20 mM imidazole, 0.5 mM DDM and 10% glycerol; and then 40 column volumes of 50 mM Tris-HCl pH 8.0, 300 mM NaCl, 40 mM imidazole, 0.5 mM DDM and 10% glycerol to remove non-specifically bound proteins. Target proteins were eluted with 50 mM Tris pH 8.0, 300 mM NaCl, 250 mM imidazole, 0.5 mM DDM and 10% glycerol in five fractions of 2 mL volume each. Eluted fractions containing the target proteins were pooled and concentrated to 0.5 mL using an Amicon Ultra-15 Centrifugal Filter Unit concentrator (MWCO 100 kDa, UFC910024) before separation on the Superose 6 Increase 10/300 column (GE Life Sciences) equilibrated with Tris (20 mM Tris-HCl pH 8.0, 150 mM NaCl, 0.5 mM DDM and 10% glycerol) size exclusion buffer. Size exclusion chromatography (SEC) shows three peaks eluting off the column: the first peak at ∼8.6 mL, the second peak at ∼12.6 mL (near the 670 kDa standard), and the third peak at ∼15.6 mL (near the 158 kDa standard) (**Fig. S5A**). While negative stain EM of the first peak displays protein aggregation, sample from the second peak displays distinct particles in what appears to be the formation of a larger complex. Based upon SDS-PAGE analysis, the ∼12.6 mL peak appears to yield 3 bands of roughly equal intensity when stained with Coomassie (Fig. S5A), which may indicate that not all 4 subunits co-purify under our conditions. Single-particle cryo-EM and 2D classification of the ∼12.6 mL peak result in barbell-like particles (**Fig. S5B**).

### cryo-EM sample preparation and data collection for complex isolated from cells over-expressing PA3211-PA3214

The fraction containing a high molecular weight complex containing His-tagged PA3211 and any co-purifying PA3212-PA3214 subunits was concentrated to 0.23 mg/mL using the Amicon Ultra-0.5 centrifugal filter unit concentrator (MWCO 100 kDa, UFC510096). Continuous carbon grids (Quantifoil R 2/2 on Cu 300 mesh grids + 2 nm Carbon, Quantifoil Micro Tools, C2-C16nCu30-01) were glow-discharged for 5 s in a PELCO easiGlow Glow Discharge Cleaning System (Ted Pella). 2.5 μL of freshly prepared sample was added to the glow-discharged grids, which were prepared using a Vitrobot Mark IV (Thermo Fisher Scientific). Grids were blotted with a blot force of 2 for 3 s at 4°C with 100% chamber humidity and then plunge-frozen into liquid ethane. Grids were clipped for data acquisition and screened at the NYU cryo-EM core facility on the Talos Arctica (Thermo Fisher Scientific) equipped with a K3 camera (Gatan). The grids were selected for data collection on the basis of ice quality and particle distribution. The selected cryo-EM grid was imaged on two separate sessions at NYSBC on Krios-1, a Titan Krios G3 electron microscope (Thermo Fisher Scientific) equipped with a K3 direct electron detector with a BioContinuum energy filter (Gatan). Super-resolution movies were collected at 300 kV using Leginon at a nominal magnification of 81,000x, corresponding to a nominal pixel size of 1.083 Å. 6,057 movies were collected at 0° tilt over a defocus range of −0.8 to −3.5 µm and each movie consisted of 30 frames with a total dose of 51.43 e-/Å2. 3,929 additional movies were collected at 30° tilt over a defocus range of −0.8 to −3.5 µm and each movie consisted of 30 frames with a total dose of 51.51 e-/Å2. Further data collection parameters are shown in **Table S2**.

### Cryo-EM data processing of the full-length complex sample

6,057 un-tilted and 3,929 tilted movies were first aligned with Relion (67), using the in-built implementation of motion correction, then transferred to CryoSPARC (68) for curation and patch-CTF estimation (Fig. S6A). 4,242,449 particles picked with WARP (76) were subjected to three rounds of 2D classification (**Fig. S6B**). An ab-initio model was generated using the remaining 1,796,259 particles, followed by non-uniform refinement. Seven rounds of 3D heterogeneous refinement were used to remove junk particles and the remaining 545,271 particles were subjected to three rounds of non-uniform refinement, followed by 2 local refinements with a mask around PA3214.

We then used these extensively cleaned particles as templates to re-pick particles using Topaz (70) (**Fig. S6C**). 6,814,327 particles were extracted using a raw box size of 256 pixels (binned 4x), and split into 8 groups; each group was then subjected to one round of 2D classification. The particles selected from the resulting 2D classification jobs were then independently classified via ab-initio modelling followed by 3D heterogeneous refinement. The remaining 3,010,123 particles were re-extracted with a binning of 2x, and subjected to further cleaning with 3 rounds of 3D heterogeneous refinements alternated with rounds of 2D classifications. Each round of 3D classification had at least 1 decoy model which was generated using ab-initio job from junk particles rejected by the initial 2D classification job. The 1,574,468 particles from the final rounds were finally refined via 3D homogeneous refinement to 4.5 Å. At this stage, the box size used allowed us to visualize PA3214 with a portion of the PA3213 needle structure, which appeared fuzzy and less well defined. Two different paths were then followed from this first consensus model: one path focused on improving the resolution of PA3214 (**Fig. S6D,E**), and the other focused on particles which were re-extracted with a larger box size to investigate the full size protein and the MCE domain (**Fig. S6F,G** detailed later).

A mask around PA3214 was made to perform 3D classification into 25 classes, limiting the resolution of the alignment to 7Å, using the PCA initialization mode with 2,000 particles, and a similarity factor of 0.1 (**Fig. S6D**). Each class was then refined via 3D homogeneous refinement, leading to reconstruction from 4.5 Å (close to Nyquist limit given our 4x binning) and 12 Å. Those reconstructions fell into 3 groups: 5 higher resolution classes (378,252 particles) in which the PA3214 ring was in a closed conformation were assigned to group 1; 3 higher resolution classes (187,311) in an open conformation were assigned to group 2; group 3 was composed of the remaining 17 classes (1,023,453 particles) which corresponded to other conformations but for which refinement did not yield reconstructions in which density was well defined enough to unambiguously assign domains, secondary structure, or an overall conformation. Combining these classes or further classification of these classes did not lead to more interpretable maps (**Fig. S6I**).

After re-extracting the particles without binning, particles from group 1 were run through a round of homogeneous and non-uniform refinement without symmetry (C1), leading to a ∼3 Å structure (**Fig. S6E**). This was followed by another round of 3D non-uniform refinement with a C8 symmetry, leading to a final model with an average resolution of 2.7 Å (Map 2, **Table S2, Fig. S7B**). Particles from group 2 were run through a homogeneous refinement followed by a non-uniform refinement in C1 symmetry, leading to the open conformation at an average resolution of 3.7 Å (Map 3, **Table S2, Fig. S7C**). Refinements of group 3 led to lower resolution and more featureless models. Integration of particles from group 3 into groups 1 and 2 did not improve the resolution of either group. Therefore, we did not pursue particles from group 3 further.

On the OM-facing surface of PA3214, densities consistent with the C-terminal residues of PA3213 were identified. These densities were observed in all 8 sites of PA3214, although we expect PA3213 to be a hexamer, and therefore expected to see only six corresponding densities. Classification based solely on the C-terminal interacting residues (only ∼7 residues) was not feasible, given the very small size of this region, and the fact that binding does not lead to a larger conformational change. To address the occupancy of the PA3213 C-terminal peptide bound to PA3214, we first performed symmetry expansion followed by local refinement and 3D classifications. Several strategies were explored (including different masks, filters and target resolutions, various number of classes, initialization modes, hard vs soft classification), many leading to either similar classes, or cases where densities near the binding regions were ambiguous. The best classification was obtained using a local mask around the binding site which extends towards the center (near the location of PA3213 needle), using hard classification and PCA mode initialization. Though some classes have ambiguous densities, two classes could be more confidently assigned as being occupied by the PA3213 C-terminus (**Fig. S6E**, bottom right, class 1 and 5), or unoccupied by PA3213 (**Fig. S6E**, bottom right, class 2). These results suggest that some PA3214 monomers may indeed be unoccupied by PA3213, as expected. However, we were not able to confidently compute the proportion of PA3213 C-terminus bound to PA3214, or define the relative position/ distribution of PA3213 C-terminal tails in the context of the whole octameric PA3214 ring.

The needle portion of PA3213 within the pore of PA3214 can be seen at a lower threshold and at much lower resolution, even when global homogeneous or non-uniform refinements are used, and remains difficult to refine to higher resolution even after signal subtraction of the PA3214 ring. Attempts to refine this region of PA3213 using various masks or classification strategies did not lead to high resolution results. Our data processing pipeline did not yield a class in which PA3213 is absent, but it is possible that PA3213 may be missing from some of the particles. It is more likely that the orientation of PA3213 is variable relative to PA3214, possibly rotating within the PA3214 pore. Analysis using 3D variability tools in CryoSPARC or multi-body refinement suggests that the PA3213 needle can rotate, tilt or move up/down relative to the PA3214 ring.

The particles from the first consensus model refined with a 256 pixels box size (∼277 Å) were also re-extracted with a larger box size of 528 pixels (∼572 Å, first binned 4x) that would include the full complex within the field of view (**Fig. S6F**), leading to 1,347,391 particles (those too close to the edge being automatically removed). A first round of 2D classification was done and the 602,344 left were used for ab-initio modelling using 2 classes followed by 3D heterogeneous refinement. The 326,143 particles from the best class, which included the full complex, were then re-centered, re-extracted (binning 1.3x) and subject to a last round of 3D homogeneous refinement to an average resolution of ∼12 Å (Map 4, **Table S2, Fig. S7A**). To assess what is present at the inner membrane, we performed a homogenous refinement of the best 3D class above using the full 1,346,391 particle set (**Fig. S6G**). After re-centering the full reconstruction around the IM complex, particles were re-extracted with a box size of 280 pixels (∼303 Å, binned 2x during the analysis), and fed into two rounds of 2D classification. A round of 3D heterogenous refinement with six classes was performed on the remaining 558,000 particles. After removing the junk classes, the remaining 96,842 particles were transferred to Relion 5.0 (67) where a new round of 2D classification was performed. Two new ab-initio models were generated with the remaining particles (63,430), and the best of them used for a new round of 3D classification with 3 classes. Class 1 and 2 were finally both refined to 17 Å. None of the classes focused on the IM showed clearly defined protein density.

### Orientation of PA3214 relative to the whole complex

Due to the modest resolution of the PA3213 needle through the PA3214 ring, we carried out additional analyses to confirm the relative orientation of PA3214 with respect to PA3213 (**Fig. S6H**). We first further refined the consensus model of the full complex through rounds of ab-initio refinement followed by heterogeneous classifications. The best models (309,480 particles) were subjected to a first round of homogeneous refinement where we can see an asymmetrical barbell with one density corresponding clearly to PA3214. Two rounds of focused local refinement with a large, and then a tighter mask around PA3214 were done to improve the resolution without risk of flipping its position relative to the IM domain. Another round of 3D classification without alignment was needed to further clean the sample and local refinement of one of those classes (67,289 particles) lead to a ∼4.5 Å resolution of PA3214. At a lower threshold, the relative position of the full complex with the MCE domains could be unambiguously defined, and at higher threshold, PA3214 density could be unambiguously defined. This allowed us to fit in the PA3214 model and confirm that the N-terminus of PA3214 faces the OM, and that the needle of PA3213 emerges from the opposite side of PA3214 (**Fig. S8**).

### Model building of closed and open conformations of PA3214

We first generated an AlphaFold 2 (77) prediction, and we rigid-body fit this predicted model into the C1 map of the closed conformation using Chimera (73). Each residue was then inspected in COOT (75) and manually adjusted as needed. Density for residues 33-210 was well defined, and we built in these residues while trimming to remove residues at either terminus. We used partial output from ModelAngelo (78) to remodel an outward-facing loop (residues 42-52) which has a lower local resolution. Densities corresponding to the last 7 residues of the C-terminus of PA3213 were built de novo in COOT.

The model was then refined in PHENIX using real_space_refine (74). We iterated several times between manual adjustments in COOT and real-space refinement in PHENIX. The model was then fit into the map with C8 symmetry applied (Map 2) and, we proceeded with another few iterations of COOT adjustments and PHENIX refinements.

To build the model of the open conformation, we performed rigid-body docking of each of the eight chains individually in Chimera (73), using a monomer of the closed conformation as a starting model. Each chain was visually inspected in COOT before a final refinement in PHENIX using the rigid_body option. Because of the lower resolution of the monomers near the opening of the ring, no further refinement was performed for chains A, G and H, while chains C to F were locally refined. First, the chains were adjusted using the ISOLDE (79) implementation within ChimeraX (80), and real-space refinement was carried out on the resulting model in Phenix using ADP and minimization_global options. Although an outward-facing loop (residues 42-52) was much lower in resolution and side-chains were not unambiguously defined, we decided to not trim anything, relying on the model obtained from the higher resolution closed conformation map. **Fig. S10** shows examples of the quality of density in different regions of the models.

### Alphafold predictions of PA3211-PA3214 complex and PqiBC complex

We utilized the AlphaFold 3 prediction software available on Google DeepMind’s AlphaFold Server (55) (https://alphafoldserver.com/) to make predictions for the PA3211-PA3214 complex and the PqiBC complex. Based on prior knowledge of MCE systems (13, 16) and our structure of PA3214^sol^, we predicted a PA3211-PA3214 complex that included 2 copies of PA3211, 2 copies of PA3212, 6 copies of PA3213 and 8 copies of PA3214 (**Fig. S5C**). This was achieved by stitching together two predictions of subcomplexes to create a composite assembly. For the PA3211-PA3213 subcomplex, we used AlphaFold 3 to predict a 2:2:6 complex. For the PA3213-PA3214 subcomplex, we used the AlphaFold Server to predict a 6:8 complex, with the signal sequence of PA3214 (residues 1 - 25) excluded. To assemble the composite prediction of the PA3211-PA3214 complex, we combined the PA3211-PA3213 prediction of all of PA3211 and PA3212, and of PA3213 residues 1-257 (the predicted junction between the penultimate and final helical modules of the needle); with the PA3213-PA3214 prediction of PA3213 residues 258-312 and all of PA3214. For the PqiBC complex, we used the AlphaFold Server to predict a 6:8 complex, with the signal sequence of PqiC (residues 1 - 15) excluded. Default program parameters were used in predictions. Program outputs yielded five ranked models of protein-protein predicted interaction and predicted alignment error (PAE) plots were generated in ChimeraX (80). The PAE plots suggest that the orientation of the PA3211 and PA3212 subunits relative to one another is well predicted. The position of the MCE domains of PA3213 (N-terminus) are well-predicted relative to each other and PA3211, while the helical needle domain of PA3213 (C-terminus) is the least well predicted region (though still globally resembles the corresponding region of other MCE proteins, such as Mce1 (16)). For the prediction of the PA3213-PA3214 subcomplex, the PAE plot indicates moderate confidence in the relative position of the PA3214 subunits. However, these interactions are well supported by our experimental structures. Similarly, moderately good PAE scores are observed between PA3214 subunits and the final ∼50 residues of PA3213, corresponding to the MCE needle interacting with the pore of the PA3214 ring, and the interactions between the C-terminal peptides of PA3213 and the hydrophobic grooves on PA3214, as observed in our structure. All together, we believe our composite model faithfully represents the overall organization of the PA3211-PA3214 complex, though we note that the needle region of PA3213 varied significantly between AlphaFold predictions and likely contains significant errors in the relative positions and packing of the α-helices in any given model. In an attempt to try to better resolve the structure of the needle, we also generated 500 models of PA3213 with Alphafold 3 loaded on the Rockfish HPC cluster at Johns Hopkins University, using 100 different random seeds to probe variations in modeling. The quality of the predictions varied significantly, and the length of the needle domain also varied substantially, from ∼179 Å to ∼200 Å when measured between residues SER138 and GLY303. Overall, the iPTM range was 0.39 - 0.51 (median of 0.47), PTM range was 0.41 - 0.52 (median of 0.49) and fraction of disordered region (FDR) was 0.10 - 0.38 (median 0.23). We discarded the worst predictions for which iPTM <0.4, PTM<0.5 and FDR>0.25, and measured the lengths of the needle domains for the remaining 96 models. The length was measured between the centroid of residues R138 and the centroid of residues R303 and resulted in variations from 179.5 - 192.8 Å (median 186.9, SD=2.7). The longest prediction is still shorter than corresponding measurements made on our map, by at least ∼37 Å.

### Deep Mutational Scanning (DMS)

The workflow for DMS was adapted from previous work (17). A library of all possible single amino acid mutations in PqiC was constructed using a similar procedure as previously described (36). The library was built by oligonucleotide-directed mutagenesis, using an NNS codon (N = any base, S = G or C) to randomize each site in the open reading frame. NNS mutagenesis reactions for each codon consisted of two PCRs (from the start codon to the site of mutation, and from the site of mutation to the stop codon), and the resulting 2 PCR products for each site were fused together by overlap PCR to yield a full-length PqiC ORF with a single codon randomized (single-site library). After repeating this for each codon in PqiC, equimolar amounts of the single-site libraries were combined in a single pool that contained all mutated positions (codons 1-187). The vector (pBEL3085, a derivative of pET17b) and the pooled mutant PqiC PCR fragments were digested with SfiI and then ligated together with T4 ligase. Top10 *E. coli* cells were transformed by electroporation with the ligated product, plated on LB agar plates with carbenicillin and incubated overnight at 37°C. We obtained ∼2.86 x 10^6^ CFU, which were scraped and pooled, followed by plasmid DNA purification with the Zymo miniprep kit. The Δ*pqiC* strain (bBEL455) was transformed by electroporation with the mutant library and grown overnight at 37°C in LB media containing 200 μg/mL carbenicillin. The cultures were diluted 1:20 into fresh LB media containing 100 μg/mL carbenicillin and shaken (200 rpm) at 37°C until OD_600_ = approximately 1. The cultures were plated on either LB + 100 μg/mL carbenicillin (unselected) or LB + 100 μg/mL + 0.7% LSB (selected) plates. Colonies from each condition were separately scraped and pooled, and plasmids were extracted. For NGS sequencing, we amplified the regions corresponding to codons 1-93 (“Sequencing Library A”, oligos 5′-actttaagaaggagatatacat-3′ and 5′-ggcagttgcgtactcaggtt-3) and codons 94-187 (“Sequencing Library B”, oligos 5′-cgcaacaccctggttgcc-3′ and 5′-atctgctcgaggtgggca-3′) by PCR, to ensure complete coverage by paired-end sequencing (2 x 250). Following PCR, DNA fragments were pooled in equimolar amounts to generate the unselected and selected samples for amplicon sequencing with Illumina MiSeq 2 x 250 paired-end sequencing (NEBNext Ultra II Library Prep kit (New England Biologs #E7645) used to generate libraries for sequencing). Two biological replicates of this experiment were carried out, starting from independent transformations of the pooled plasmid library into the Δ*pqiC* strain (bBEL455). These independent transformations and all subsequent selection, sequencing, and analysis steps were carried out several months apart. Because the plasmid libraries corresponding to codons 1-93 and codons 94-187 were not cloned separately, PCR fragments from Sequencing Library A with an apparent WT sequence may be derived from truly WT plasmids, or may instead be derived from a plasmid with a mutation in codons 94-187 (the region covered by Sequencing Library B). Similarly, PCR fragments from Sequencing Library B with an apparent WT sequence may be derived from WT plasmids, or from a plasmid with a mutation in codons 1-93 (the region covered by Sequencing Library A). The effect of this is a possible over-estimation of the frequency of the WT sequence, relative to all mutants. However, this is not expected to skew the relative fitness of individual mutants relative to one another.

The data analysis of the pair-end reads was performed in a similar manner as previously described (17). Illumina MiSeq paired end reads of each replicate were first aligned to the wild type (WT) reference sequence of PqiC using bowtie2 (81). After converting the files to BAM format, samtools (82) was used to keep only properly paired reads and filter out reads with a poor mapping quality (MAPQ<42). Following a conversion to fastq format via bedtools (83), overlapping reads were merged using pandaseq (84). Primer sequences were then removed using cutadapt (85) before running python scripts previously developed (36) to remove ambiguous nucleotides, filter by length, assess correct orientation to the proper reading frame, translate to the corresponding protein sequence, and count single amino acid variants. These resulting normalized variant counts obtained for the selected and non-selected populations were used to compute the relative fitness value (𝛥*E*^*X*^_*i*_) (17). Using the fitness cost, a tolerance score (TS) (59) was calculated (using a Z-score threshold of <-3.5 and >3.5) for each residue based on the number and types of amino acid substitutions that were tolerated at that position, and derived from the Zvelebil similarity score that incorporates global amino acid distinctions (60, 61). A Tolerance Score [TS] < 0.9 was used as a threshold, as this cutoff excluded mutation tolerant residues (∼60% of all PqiC residues), combined with manually excluding residues from the signal peptide and hydrophobic core (∼20% of all PqiC residues), that led to the final functional categorization of 39 residues (∼20% of all PqiC residues). The square of the Pearson correlation coefficient (r^2^) between the two replicates is r^2^=0.852 and r^2^=0.811 on the fitness matrices of LibA and LibB respectively. Aggregation of the replicates was performed at the mutation fitness level doing an average between the two matrices (or, for cells of the matrix with an incomplete coverage, only the value of the other replicate was included). We generated a semi-automated pipeline for DMS data processing and used this for processing these data. The code and processing pipeline used for the DMS sequencing reads are available on GitHub https://github.com/ncoudray/Fitness-Mapper.

### Expression and Purification for PA3214^Δintseg^

Plasmid pBEL3448, which encodes for N-terminal 6xHis2xQH-TEV tagged PA3214^Δintseg^ (PA3214 residues 55-214) was transformed in Rosetta 2(DE3) cells (Novagen #714003) (**Table S3**). For protein expression, an overnight culture (LB + 100 µg/mL carbenicillin + 38 µg/mL chloramphenicol) was diluted 1:50 in LB (Difco) supplemented with carbenicillin (100 µg/mL) and chloramphenicol (38 µg/mL), grown at grown at 37°C with shaking to an OD600 of 0.6-0.8, and induced by the addition of L-arabinose (final concentration 0.2%). Cultures were then incubated at 37°C with shaking for 4 hours, followed by harvesting by centrifugation at 4,500g for 20 minutes. Pellets were resuspended in 50 mM Tris pH 8.0, 300 mM NaCl, 10mM Imidazole, and stored at −80°C. Cells were lysed by three passes through an Emulsiflex-C3 cell disruptor (Avestin), then centrifuged at 32,000g for 35 minutes at 4°C to pellet cell debris. The supernatant was incubated with Ni Sepharose resin (Cytiva 17-5318-02) for 30 minutes at 4°C on a rocker, and then washed with 10 column volumes of 50 mM Tris-HCl pH 8.0, 300 mM NaCl, 10 mM imidazole; then 20 column volumes of 50 mM Tris-HCl pH 8.0, 300 mM NaCl, 20 mM imidazole; and then 40 column volumes of 50 mM Tris-HCl pH 8.0, 300 mM NaCl, 40 mM imidazole to remove non-specifically bound proteins. Target protein was eluted with 10 mL 50 mM Tris pH 8.0, 300 mM NaCl, 250 mM imidazole in five fractions of 2 mL volume each. Eluted fractions containing target proteins were pooled and concentrated to 0.5 mL using an Amicon Ultra-15 Centrifugal Filter Unit concentrator (MWCO 10 kDa, UFC901024) before separation on the Superdex 200 Increase 10/300 column (Cytiva) equilibrated Tris (20 mM Tris-HCl pH 8.0, 150 mM NaCl, 0.5 mM DDM and 10% glycerol) size exclusion buffer. Fractions containing PA3214^Δintseg^ peak 1 and 2 (Fig. S9C) were collected and applied to grids for negative stain EM.

### Negative stain EM

To prepare grids for negative-stain EM analysis (**Fig. S9D**), fresh samples of protein after size exclusion chromatography (PA3214^sol^ and PA3214^Δintseg^ peak 1 and 2) were applied to a carbon-coated 400 mesh copper grid (Ted Pella, 01754-F), freshly glow discharged for 30 seconds. The samples were then blotted off on filter paper (Whatman 1). Immediately after blotting, 3 µL of a 2% uranyl formate solution was applied for staining and blotted off on filter paper (Whatman 1). Application and blotting of stain was repeated four times. The sample was allowed to air dry before imaging. Data were collected on the Talos F200C TEM (FEI) equipped with the Ceta-D camera at a nominal magnification of 73,000x corresponding to a pixel size of 2.04 Å /px on the sample and a defocus range of −1 to −2.18 μm defocus. Micrographs were imported into CryoSPARC and 27,059 particles were picked using automated blob picking. Particles were extracted with a box size of 128 pixels and duplicated particles were removed. 2D classification with 25,370 particles was performed using default parameters. As PA3214^Δintseg^ does not oligomerize to form an octameric ring, micrographs for this dataset did not contain particles of sufficient size to pick or process further.

### Pull-down assays

Plasmids pBEL1282 (WT *E. coli* PqiABC with an N-terminal 6xHis2xQH-TEV tag on PqiB) and pBEL3438 - pBEL3445 (6 mutant constructs derived from pBEL1282) were each separately transformed into Rosetta 2(DE3) cells (Novagen #714003) (**Table S3**). Plasmids pBEL3081 (WT *P. aeruginosa* PA3211-PA3214 with a C-terminal 2xHQ6xHis tag on PA3214), pBEL3447 and pBEL3449-56 (8 mutant constructs), as well as an untagged control construct (pBEL3446), were each separately transformed in Rosetta 2(DE3) cells (Novagen #714003) (**Table S3**). Due to the inherent stickiness of PA3214 to NiNTA beads, the His-tag on PA3211-PA3214 constructs was placed on PA3214 as the bait protein. The *pqiB, pqiC, PA3213* and *PA3214* regions were mutated using Gibson assembly. Whole plasmid sequencing was performed by Plasmidsaurus to confirm all mutant plasmids.

For protein expression, an overnight culture (LB + 100 µg/mL carbenicillin + 38 µg/mL chloramphenicol) was diluted 1:40 in LB (Difco) supplemented with carbenicillin (100 µg/mL) and chloramphenicol (38 µg/mL), grown at 37°C with shaking to an OD600 of 0.6-0.8, and induced by the addition of L-arabinose (final concentration 0.2%). Cultures were then incubated at 37°C with shaking for 4 hours, followed by harvesting by centrifugation at 4,500g for 20 minutes. Pellets were resuspended in 50 mM Tris pH 8.0, 300 mM NaCl, 10% glycerol, 10mM imidazole, and stored at −80°C. Cells were lysed by three passes through an Emulsiflex-C3 cell disruptor (Avestin), then centrifuged at 32,000g for 35 minutes at 4°C to pellet cell debris. The clarified lysate underwent ultracentrifugation at 37,000 rpm (182,460g) for 49 minutes at 4°C Fiberlite F37L-8 x 100 Fixed-Angle Rotor (Thermo Fisher Scientific, 096-087056). The supernatant was discarded and the membrane fraction was solubilized in 50 mM Tris pH 8.0, 300 mM NaCl, 10% glycerol, 25 mM n-DodecylB-D-maltoside (DDM) by rocking overnight at 4°C. Insoluble debris were pelleted by ultracentrifugation at 37,000 rpm for 49 minutes at 4°C. Solubilized membranes were incubated with Ni Sepharose resin (Cytiva 17-5318-02) for 30 minutes at 4°C on a rocker, and then washed with ∼160 column volumes of 50 mM Tris-HCl pH 8.0, 300 mM NaCl, 40 mM imidazole, 0.5 mM DDM and 10% glycerol to remove non-specifically bound proteins. Target proteins were eluted with 50 mM Tris pH 8.0, 300 mM NaCl, 250 mM imidazole, 0.5 mM DDM and 10% glycerol in five fractions of 2 mL volume each. Eluted fractions containing target protein were pooled and mixed with 5x SDS-PAGE loading buffer.

### Generation of PqiB, PqiC, PA3213 and PA3214 polyclonal antibodies

Custom antibodies were generated by Capra Sciences, either against a peptide epitope (PA3213) or against protein (PA3214, PqiB and PqiC).

Protein preparation for antibody generation was carried out as follows. Plasmids pBEL1163 (N-terminal 6xHis-TEV tagged PqiB(39-546)), pBEL1296 (PelB-6xHis-TEV-PqiC(15-187), pBEL2277 (6xHis2xQH-TEV-PA3214(27-214), were each separately transformed into Rosetta 2(DE3) cells (Novagen #714003) (**Table S3**). For protein expression, an overnight culture (LB + 100 µg/mL carbenicillin + 38 µg/mL chloramphenicol) was diluted 1:50 in LB (Difco) supplemented with carbenicillin (100 µg/mL) and chloramphenicol (38 µg/mL), grown at grown at 37°C with shaking to an OD600 of 0.6-0.8, and induced by the addition of IPTG (final concentration 1 mM IPTG) for pBEL1163 and pBEL1296 or arabinose (final concentration 0.2%) for pBEL2277. Cultures were then incubated at 37°C with shaking for 4 hours, followed by harvesting by centrifugation at 4,500g for 20 minutes. Pellets were resuspended in 50 mM Tris pH 8.0, 300 mM NaCl, flash frozen in liquid nitrogen, and stored at −80°C. Cells were lysed by three passes through an Emulsiflex-C3 cell disruptor (Avestin), then centrifuged at 32,000g for 35 minutes at 4°C to pellet cell debris. The supernatants were incubated with Ni Sepharose resin (Cytiva 17-5318-02) for 30 minutes at 4°C on a rocker, and then washed with 10 column volumes of 20 mM Tris-HCl pH 8.0, 150 mM NaCl, 10 mM imidazole; then 20 column volumes of 20 mM Tris-HCl pH 8.0, 150 mM NaCl, 20 mM imidazole; and then 40 column volumes of 20 mM Tris-HCl pH 8.0, 150 mM NaCl, 40 mM imidazole to remove non-specifically bound proteins. Target proteins were eluted with 10 mL 50 mM Tris pH 8.0, 300 mM NaCl, 250 mM imidazole in five fractions of 2 mL volume each. Eluted fractions containing target proteins were pooled and concentrated to 0.5 mL using an Amicon Ultra-0.5 Centrifugal Filter Unit concentrator (MWCO 100 kDa, UFC510096) before separation on the Superdex 200 Increase 10/300 column (Cytiva) equilibrated with Tris (20 mM Tris-HCl pH 8.0, 150 mM NaCl) size exclusion buffer. The peak containing our protein of interest was pooled and sent to Capra Sciences for antibody generation. To acquire specific chicken anti-PqiB and anti-PqiC antibodies, the egg yolk from 8 eggs (per antigen) were used to generate an affinity purified polyclonal antibody, using a column coupled with antigen (AKTA HPLC). Due to the low yield of PA3213, a synthetic peptide (PA3213^115-129^: SGTPASPMLEGKDGK) was designed and synthesized by Capra Science for antibody generation. The specificity of PqiB, PqiC, PA3213 and PA3214 antibodies were validated by blotting of WT and knockout cell lysates, as well as purified protein.

### Western blotting

Samples were separated on an SDS-PAGE gel and transferred to a nitrocellulose membrane using the Trans-Blot Turbo System (Bio-Rad Laboratories). The membranes were blocked in Phosphate Buffered Saline Tween-20 (PBST, 1X PBS + 0.05% Tween-20) containing 5% milk for 1 hour at room temperature or overnight at 4°C with agitation. To probe PqiB, the membranes were incubated with primary antibody in PBST + 5% BSA, chicken polyclonal anti-PqiB at a final concentration of 1 µg/mL. To probe PqiC, the membranes were incubated with a primary antibody in PBST + 5% BSA, chicken polyclonal anti-PqiC at a final concentration of 1 µg/mL. To probe PA3213, the membranes were incubated with a primary antibody in PBST + 5% BSA, chicken polyclonal anti-PA3213 at a final concentration of 0.1 µg/mL. To probe PA3214, the membranes were incubated with a primary antibody in PBST + 5% BSA, rabbit polyclonal anti-PA3213 at a final concentration of 0.1 µg/mL. Membranes were incubated in primary antibody solution for one hour at room temperature. The membranes were then washed 3 times with PBST and incubated with Donkey anti-chicken IgG IRDye® 680RD (LI-COR Biosciences, Catalog #926-68075), Donkey anti-chicken IgG IRDye® 800CW (LI-COR Biosciences, Catalog #926-32218) or Goat anti-rabbit IgG IRDye® 800CW (LI-COR Biosciences, Catalog #926-32211) secondary antibodies in PBST + 5% BSA for 1 hour at room temperature with agitation. The membranes were then washed 3 times with PBST and imaged on a LICOR Odyssey Classic.

### Software packages

Several structural biology applications used in this project and noted above were compiled and configured by SBGrid (86). Molecular graphics and analyses were performed using UCSF ChimeraX, developed by the Resource for Biocomputing, Visualization, and Informatics at the University of California, San Francisco, with support from National Institutes of Health R01-GM129325 and the Office of Cyber Infrastructure and Computational Biology, National Institute of Allergy and Infectious Disease (80).

### Generative AI usage

Gemini and ChatGPT were used in refining the code for DMS analysis at several steps. All outputs were manually inspected by authors.

## Data Availability

The cryo-EM coordinates and associated maps have been deposited at the PDB and Electron Microscopy Data Banks for the soluble PA3214^sol^ complex (PDB: 13HQ, EMDB: EMD-77071 for map 1), the closed conformation of PA3214 (PDB: 13HR, EMDB: EMD-77072 for map 2), the open conformation of PA3214 (PDB: 13HS, EMDB: EMD-77073 for map 3), and the full complex (EMDB: EMD-77074). Cryo-EM data will be deposited to the Electron Microscopy Public Image Archive (EMPIAR). Raw sequencing reads were deposited to the Sequence Read Archive under BioSample accessions SAMN58692440 and SAMN58692441. DMS data were deposited to MaveDB with accession code urn:mavedb:00001273-a. The model of PA3211-PA3214 stitched together from two subcomplexes generated by alphafold3 is available in ModelArchive (modelarchive.org) with the accession code ma-48usm.

## Code availability

The code used to process DMS data is available on GitHub at https://github.com/ncoudray/Fitness-Mapper.

## Acknowledgments

We thank members of the Bhabha + Ekiert Labs for helpful discussions. We thank Cristina Santarossa, Benjamin Thomson, Yongbo Ding, Margot Di Cesare, Ethan Yarberry, Jaskamaljot Kaur Banwait, Paris Watson and Kacie McCarty for critical reading and feedback on our manuscript. We gratefully acknowledge the following funding sources: National Institutes of Health (NIH) grant (R35GM128777) to D.C.E., Pew Charitable Trusts grant (PEW-00033055) to G.B., National Science Foundation Graduate Research Fellowship Program grant (2021318502) to S.I.G. Cryo-EM grids were prepared and screened at the NYU Langone Health’s Cryo-Electron Microscopy Laboratory, which is partly supported by grants NIH/NCI P30CA016087, and we thank William Rice and Bing Wang for assistance with cryo-EM grid screening and microscope operation. Some of this work was performed at the Simons Electron Microscopy Center at the New York Structural Biology Center, with major support from the Simons Foundation (SF349247). Some Cryo-EM data collection was performed at the Pacific Northwest Center for Cryo-EM (PNCC), supported by NIH grant R24GM154185. For data processing, we used compute resources maintained by the Advanced Research Computing at Hopkins (ARCH) core facility (rockfish.jhu.edu), which is supported by the National Science Foundation (NSF) grant number OAC1920103. Initial data processing was carried out using the HPC cluster at NYU School of Medicine. We thank Joe Biondo, Ricardo Jacomini, Andre Barbosa, Matt Protzman and Jaime Combariza for helpful discussions regarding HPC utilization and resources. Sequencing was performed by the Genome Technology Center (RRID: SCR_017929) at NYU School of Medicine for the first DMS replicate, and by the Integrated Genomics Center (RRID SRC_018669) at Johns Hopkins Department of Genetic Medicine for the second replicate. This Genome Technology Center at NYU School of Medicine is partially supported by the Cancer Center Support Grant P30CA016087 at the Laura and Isaac Perlmutter Cancer Center.

## Author contributions

S.I.G., N.C., R.R., G.B. and D.C.E. were responsible for project conceptualization, formal analysis and validation. S.I.G., N.C. and R.R. were responsible for methodology and data curation. G.B. and D.C.E. supervised the project and provided resources. S.I.G., G.B. and D.C.E. were supported by funding. All authors edited the manuscript.

