## Supplementary Figures (S1-S10) for "Interactions of outer membrane lipoproteins *P. aeruginosa* PA3214 and *E. coli* PqiC with their MCE protein binding partners, PA3213 and PqiB"

**Supplementary Figure S1**

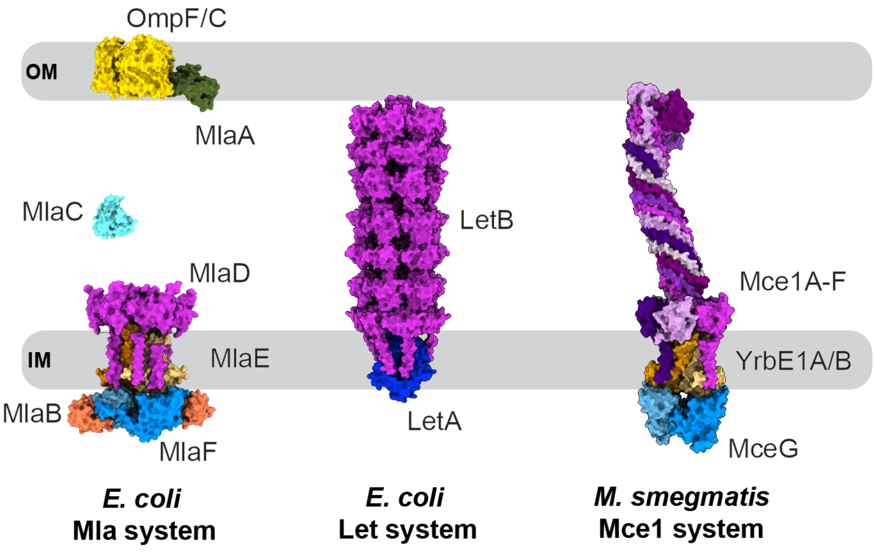


**Figure S1. Examples of MCE systems in *E. coli* and mycobacteria.** *E. coli* Mla system (PDB 6XBD, 5UWA, 5NUR), *E. coli* Let system (PDB 9N8W) and *M. smegmatis* Mce1 (PDB 8FEE) MCE systems in the context of a double-membraned cell envelope. MalD, LetB and Mce1A-F are the MCE proteins in these systems.

#### Supplementary Figure S2

A

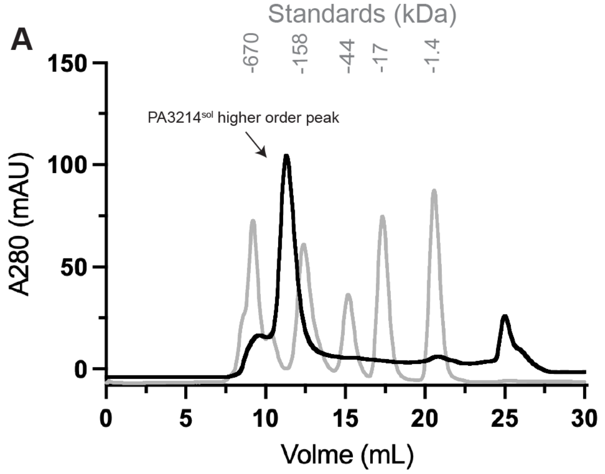


B

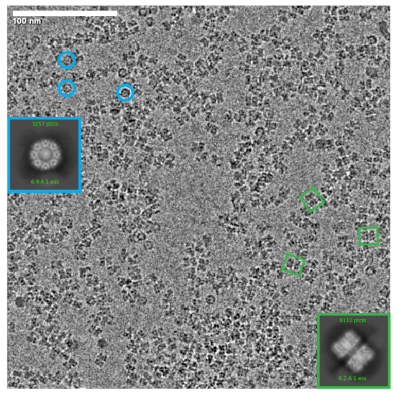


C

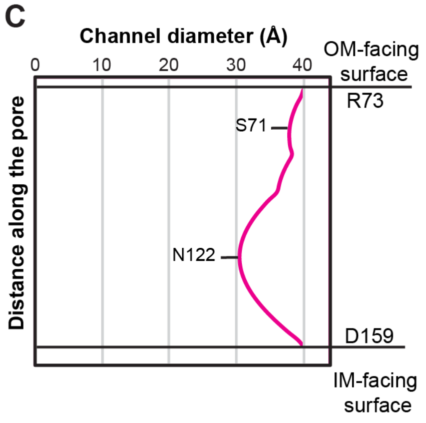


D

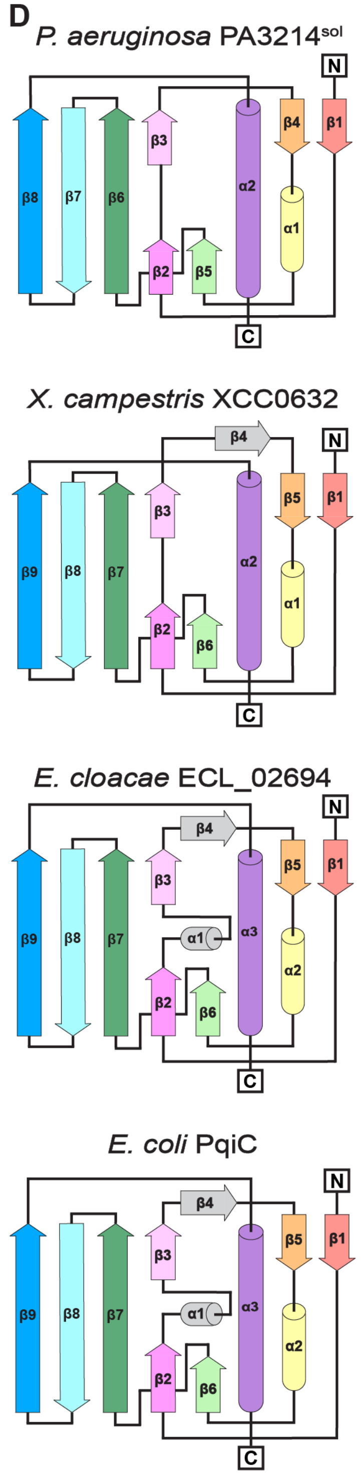


E

**Structural Similarity (E-value)**

| | *P. aeruginosa* PA3214 | *X. campestris* XCC0632 | *E. cloacae* ECL_02694 | *E. coli* PqiC |
|---|---|---|---|---|
| *X. campestris* XCC0632 | 3.31 x $10^{-18}$ | 8.58 x $10^{-35}$ | 6.77 x $10^{-10}$ | 1.92 x $10^{-9}$ |
| *E. cloacae* ECL_02694 | 2.6 x $10^{-10}$ | 1.17 x $10^{-8}$ | 4.57 x $10^{-36}$ | 2.36 x $10^{-28}$ |
| *E. coli* PqiC | 1.8 x $10^{-10}$ | 7.16 x $10^{-9}$ | 1.12 x $10^{-28}$ | 1.64 x $10^{-37}$ |

**Sequence Identity**

| | *P. aeruginosa* PA3214 | *X. campestris* XCC0632 | *E. cloacae* ECL_02694 | *E. coli* PqiC |
|---|---|---|---|---|
| *P. aeruginosa* PA3214 | 100% | 33.5% | 13.5% | 12.6% |
| *X. campestris* XCC0632 | 33.5% | 100% | 16.4% | 13.2% |
| *E. cloacae* ECL_02694 | 13.5% | 15.7% | 100% | 78.8% |
| *E. coli* PqiC | 12.6% | 13.8% | 78.8% | 100% |

F

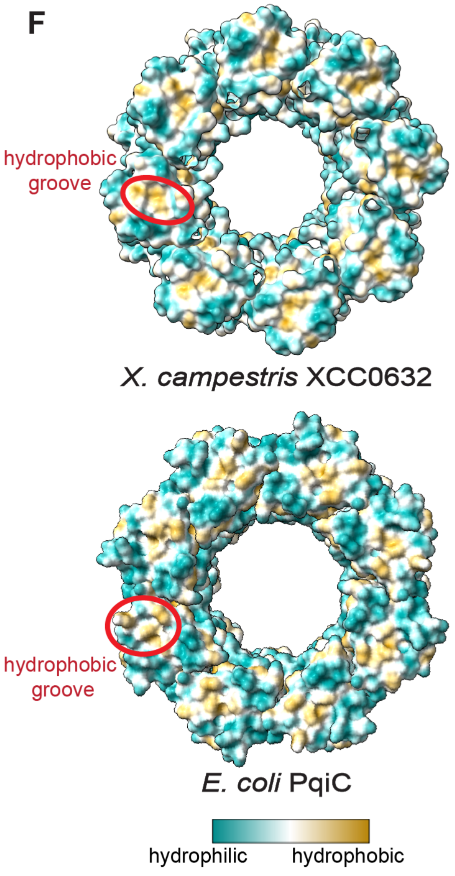


G

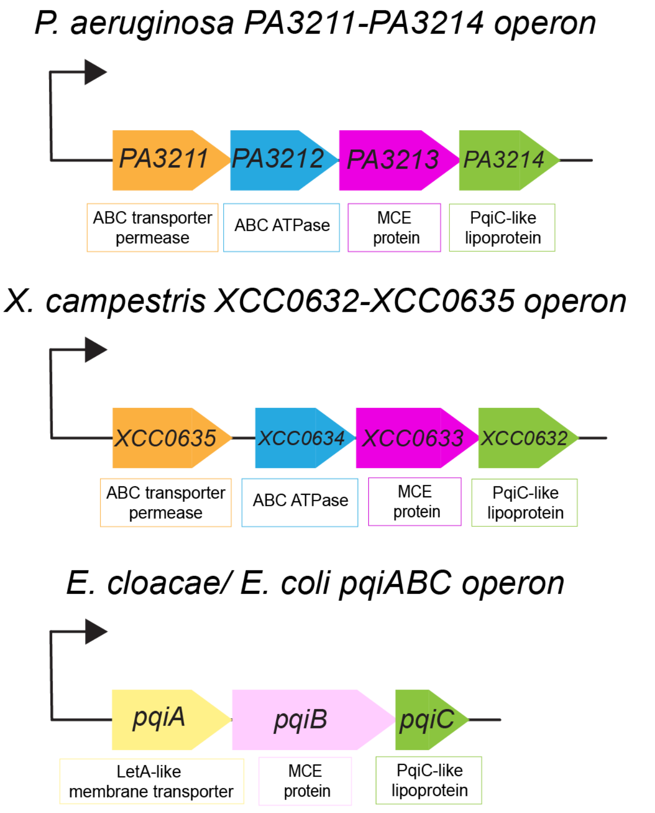

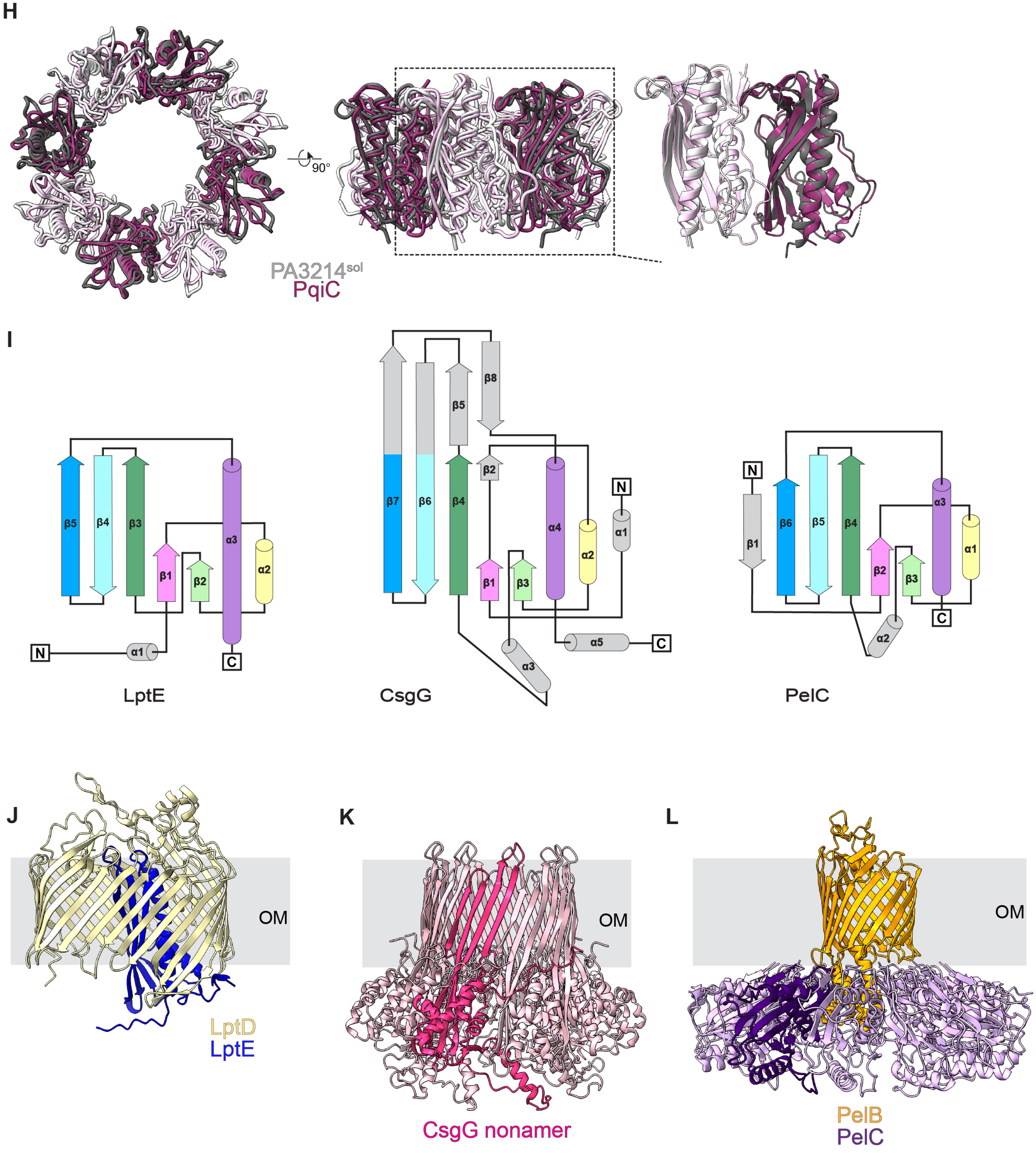
H
90°
PA3214sol
PqiC
I
β5
β4
β3
β1
β2
α3
α2
α1
N
C
LptE
β7
β6
β5
β4
β8
β2
β1
β3
α4
α2
α1
α3
α5
N
C
CsgG
N
β1
β6
β5
β4
β2
β3
α3
α1
α2
C
PelC
J
OM
LptD
LptE
K
OM
CsgG nonamer
L
OM
PelB
PelC

**Figure S2. Biochemical and structural characterization of PA3214$^{sol}$.** *A,* a representative size exclusion chromatogram for PA3214$^{sol}$ (black) overlaid with standards (gray). The PA3214$^{sol}$ higher order complex elutes between the 158 and 670 kDa protein standard. *B,* example of a typical PA3214$^{sol}$ cryo-EM micrograph. Examples of top views are indicated with blue circles and the blue inset shows a representative 2D class. Examples of side views are indicated with green rectangles and the green inset shows a representative 2D class, highlighting the non-specific interaction between two rings. *C,* diameter of the PA3214$^{sol}$ pore (magenta) along the length of the lipoprotein from the OM-facing surface to the IM-facing surface, measured using CHAP (87). *D,* topology diagrams of *P. aeruginosa* PA3214$^{sol}$, *X. campestris* XCC0632 (PDB 2IQI), *E. cloacae* PqiC (PDB 6OSX), and *E. coli* PqiC (PDB 8Q2C) highlighting structural homology compared to PA3214$^{sol}$ (colored elements) and features not found in PA3214 in gray. *E,* comparison of structural similarity and sequence identity between *P. aeruginosa* PA3214$^{sol}$, *X. campestris* XCC0632, *E. cloacae* PqiC and *E. coli* PqiC. Values obtained from FoldSeek search (44). *F,* molecular surface representation of the OM-facing surfaces of *X. campestris* XCC0632 (PDB 2IQI) and *E. coli* PqiC (PDB 8Q2C) colored by hydrophobicity. Hydrophobic groove indicated by red circle. *G,* schematic of *P. aeruginosa PA3211-PA3214*, *X. campestris XCC0632-XCC0635*, and *E. cloacae* / *E. coli pqiABC* operons. *H,* Overlay of PA3214$^{sol}$ and PqiC, aligned on the octamer. *I,* topology diagrams of LptE (PDB 4N4R), CsgG (PDB 7BRM) and PelC (5T10) highlighting structural homology compared to PA3214 (colored elements) and features not found in PA3214 in gray. *J,* cartoon representation of the LptDE complex (PDB 4N4R) relative to the OM (gray). *K,* cartoon representation of CsgG nonamer (PDB 7BRM), with a monomer highlighted in hot pink, relative to the OM (gray). *L,* a cartoon representation of the PelBC complex (PDB 9H80), with a monomer of PelC highlighted in dark purple, relative to the OM (gray).

**Supplementary Figure S3**

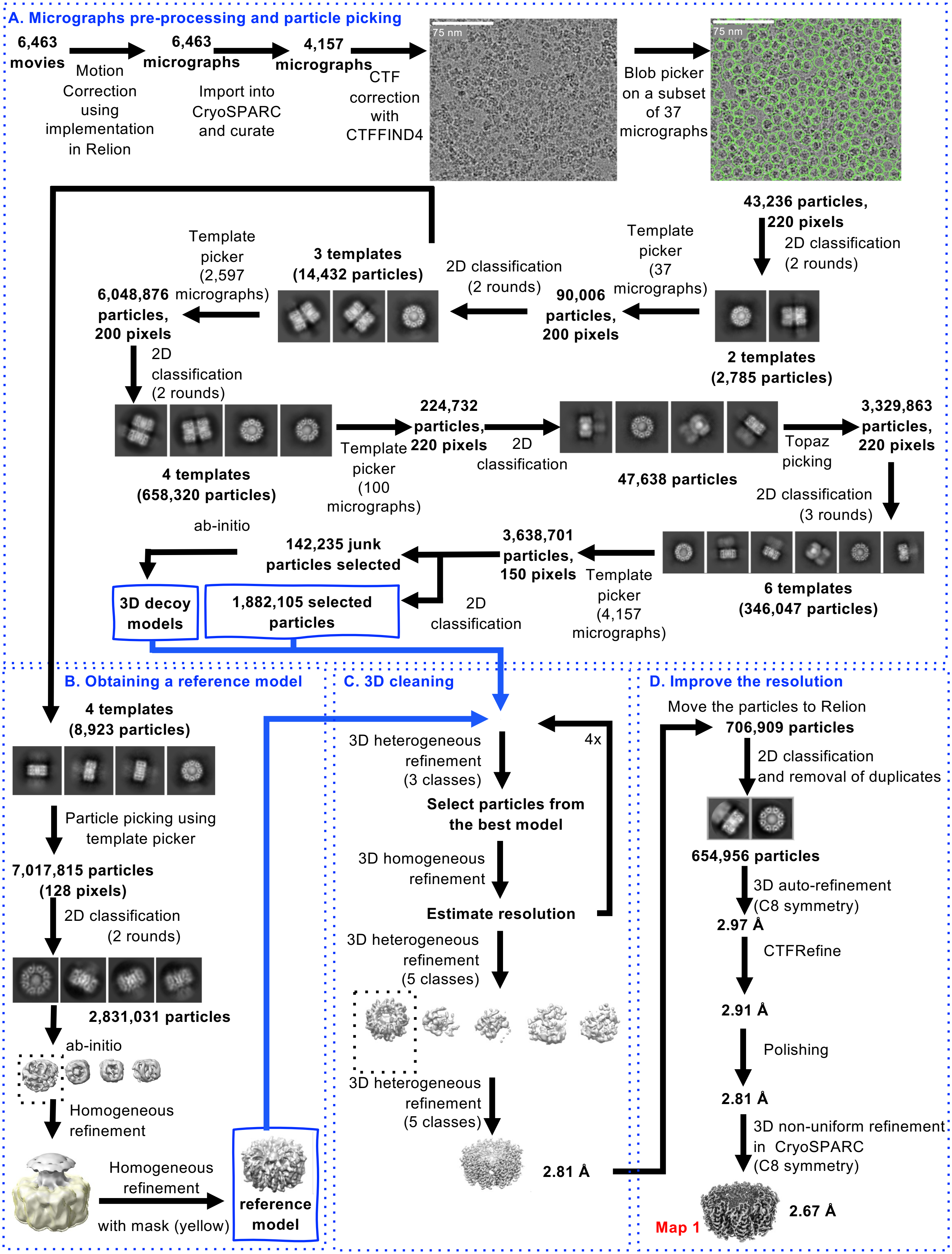

**Figure S3. Cryo-EM data processing workflow for the structure of PA3214$^{sol}$.** *A*, micrographs pre-processing and particle picking. *B*, Generating a reference model of a single PA3214$^{sol}$ ring. *C*, 3D cleaning of the dataset. *D*, steps implemented to improve the resolution, leading to the final model. See Methods for details.

### Supplementary Figure S4

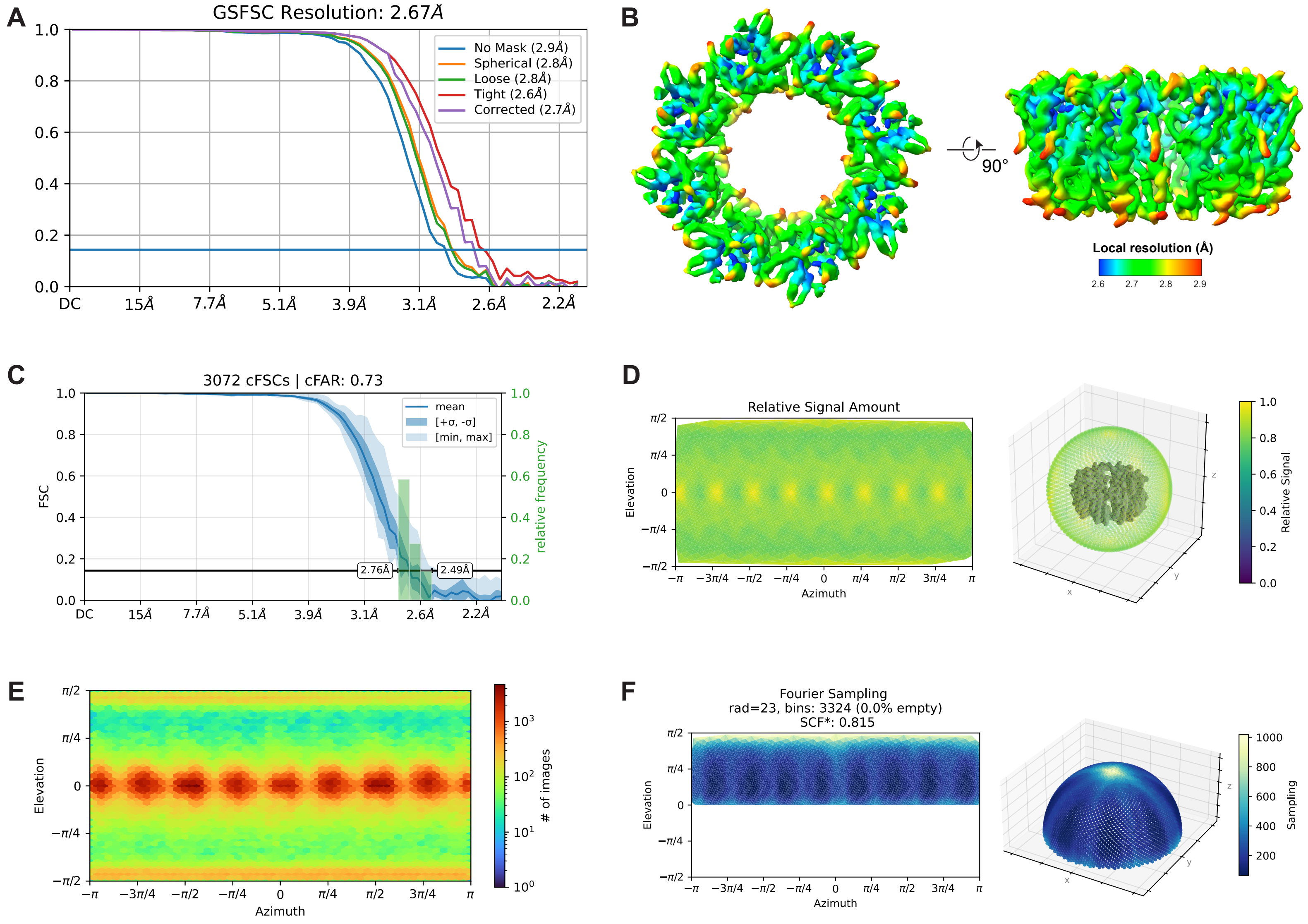


**Figure S4. Map validation corresponding to the workflow in Fig. S3 (Map 1).** *A*, overall Gold Standard Fourier Shell Coefficient (GSFSC) curve. *B*, local resolution of PA3214$^{sol}$ map. *C*, canonical FSC (cFSC) curve. *D*, relative signal amount by viewing direction plots. *E*, viewing direction distribution. *F*, fourier sampling plot. All plots were generated with CryoSPARC.

#### Supplementary Figure S5

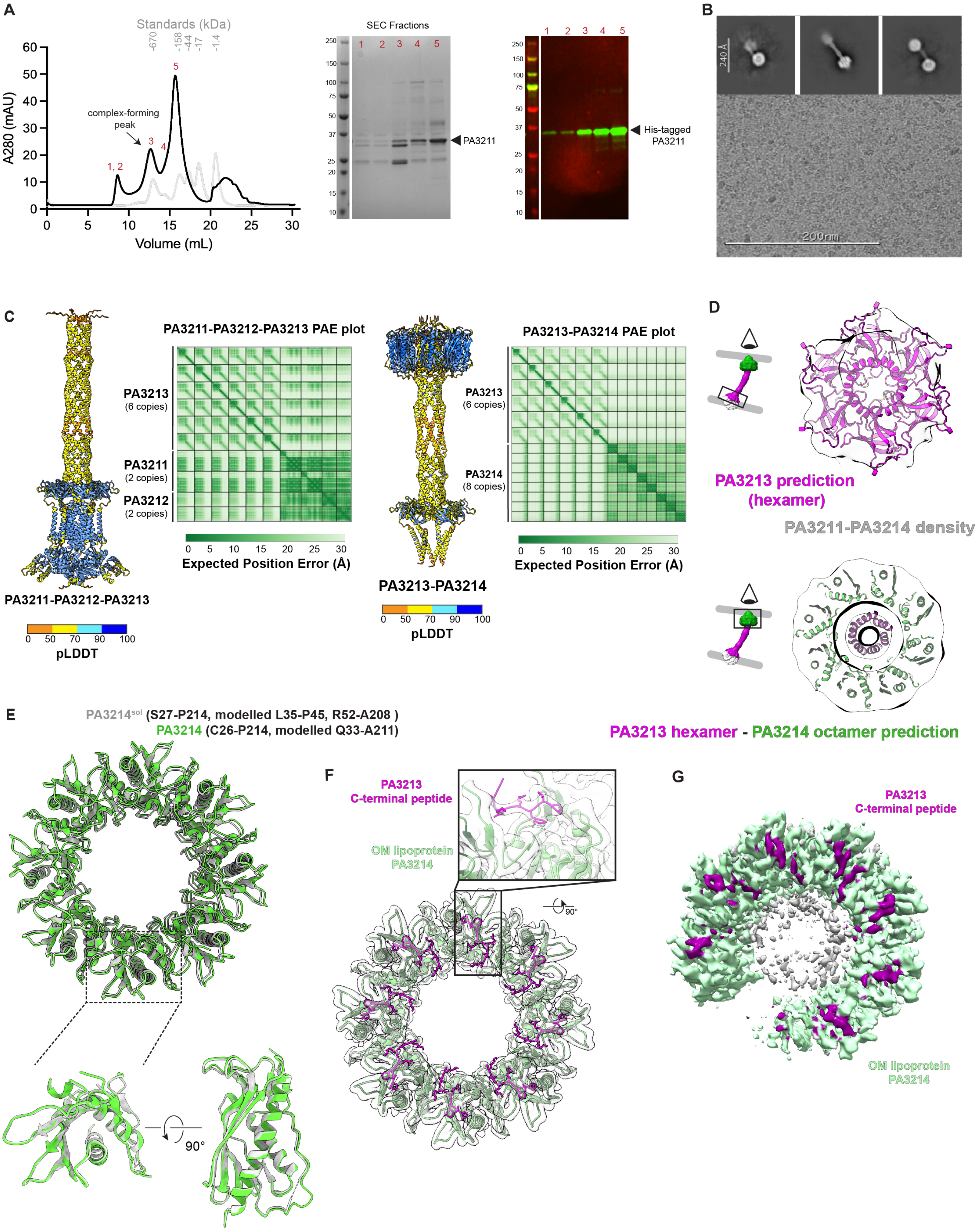

**Figure S5. Biochemical and structural characterization of the PA3211-PA3214 complex.** *A,* representative size exclusion chromatogram for PA3211-PA3214 complex (black) overlaid with standards (gray) and 5 peak fractions analyzed shown in red. Corresponding SDS-PAGE stained with coomassie, and Western blot are shown. *B,* representative cryo-EM micrograph for PA3211-PA3214 sample, with an inset showing 2D classes. *C,* AlphaFold 3 predictions of PA3211-PA3212-PA3213 and PA3213-PA3214 that were used to prepare a composite prediction of the PA3211-PA3214 complex in Fig. 2C. Confidence statistics for each prediction are shown with pLDDT (Predicted Local Distance Difference Test) values colored as in the corresponding key; predicted alignment error (PAE) values between the proteins of each prediction plotted on the corresponding heatmap. *D,* top view of AlphaFold 3 PA3211-PA3214 complex prediction showing the MCE hexameric ring assembly (magenta), or the PA3214 octamer assembly (green) docked into the low resolution consensus map. *E,* overlay of the cartoon representation of $PA3214^{sol}$ structure (gray) with the full-length PA3214 structure (green). *F,* model of PA3214 (green) and of the C-terminus of PA3213 (magenta) in the cryo-EM map (Map 2, transparent gray), with inset showing the density in that region. *G*, density for PA3214 in the open conformation (Map 3) with density corresponding to the PA3213 C-terminal peptide shown in magenta.

#### Supplementary Figure S6

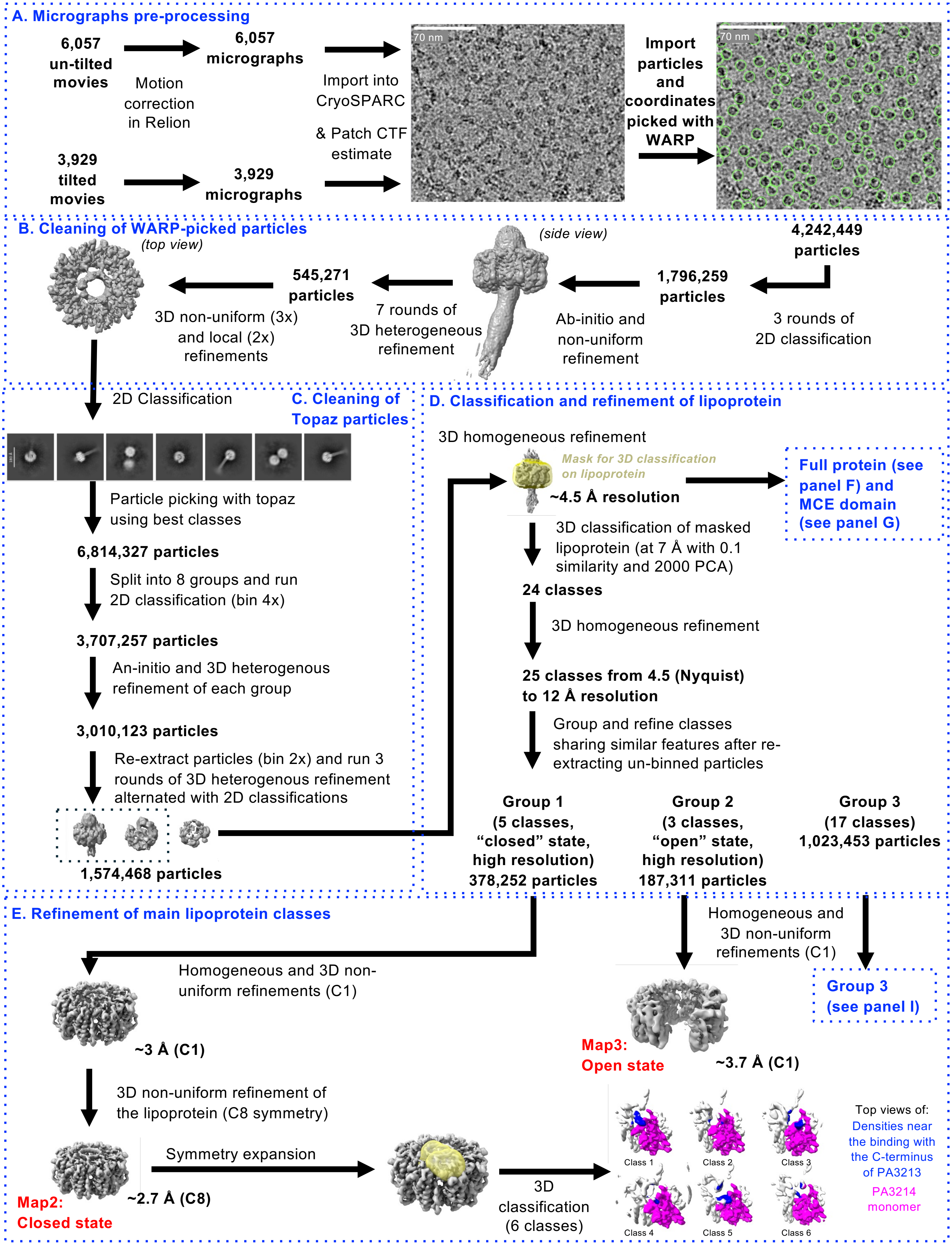

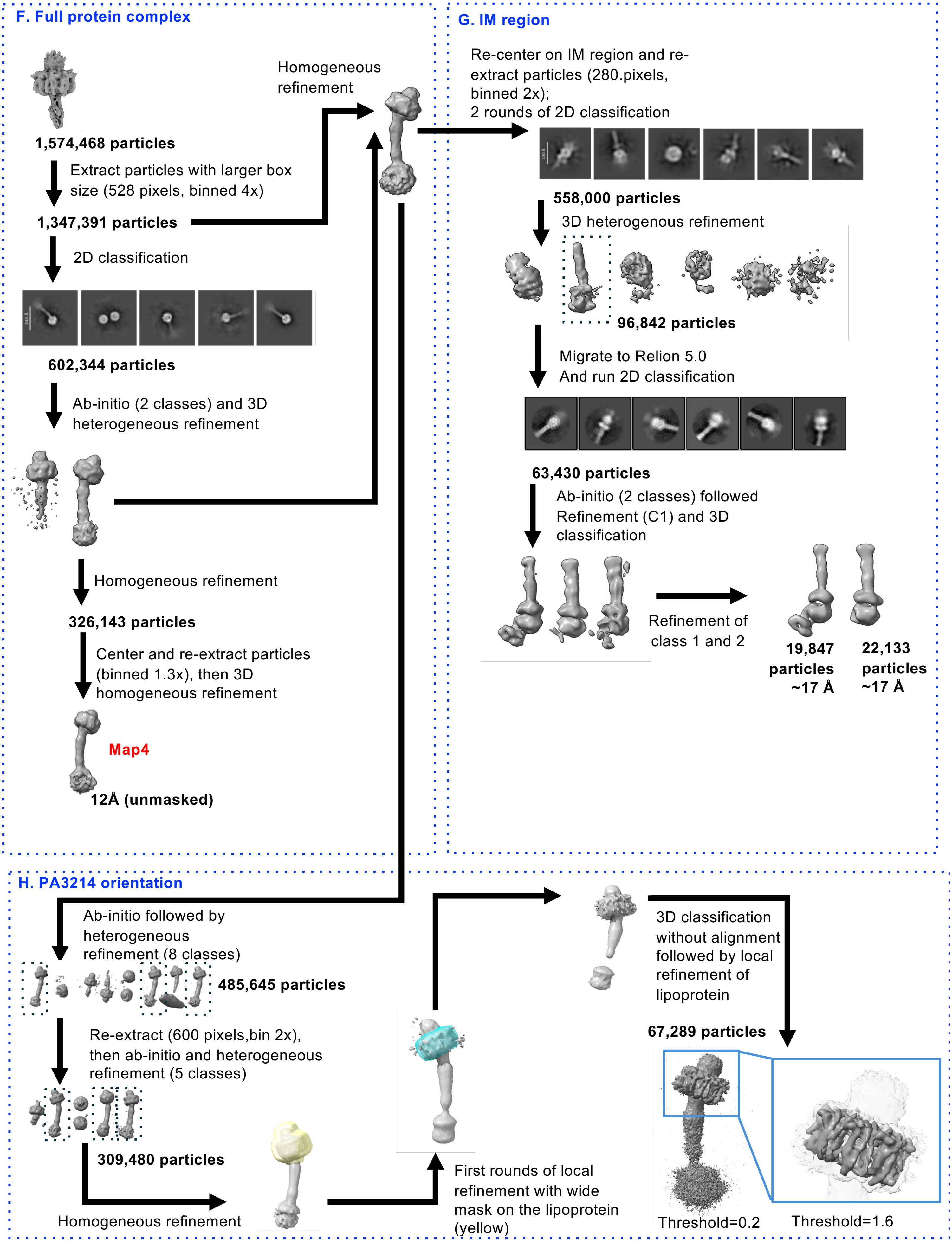

F. Full protein complex
1,574,468 particles
Extract particles with larger box size (528 pixels, binned 4x)
1,347,391 particles
Homogeneous refinement
2D classification
602,344 particles
Ab-initio (2 classes) and 3D heterogeneous refinement
Homogeneous refinement
326,143 particles
Center and re-extract particles (binned 1.3x), then 3D homogeneous refinement
Map4
12Å (unmasked)
G. IM region
Re-center on IM region and re-extract particles (280.pixels, binned 2x);
2 rounds of 2D classification
558,000 particles
3D heterogenous refinement
96,842 particles
Migrate to Relion 5.0
And run 2D classification
63,430 particles
Ab-initio (2 classes) followed Refinement (C1) and 3D classification
Refinement of class 1 and 2
19,847 particles ~17 Å
22,133 particles ~17 Å
H. PA3214 orientation
Ab-initio followed by heterogeneous refinement (8 classes)
485,645 particles
Re-extract (600 pixels,bin 2x), then ab-initio and heterogeneous refinement (5 classes)
309,480 particles
Homogeneous refinement
First rounds of local refinement with wide mask on the lipoprotein (yellow)
3D classification without alignment followed by local refinement of lipoprotein
67,289 particles
Threshold=0.2
Threshold=1.6

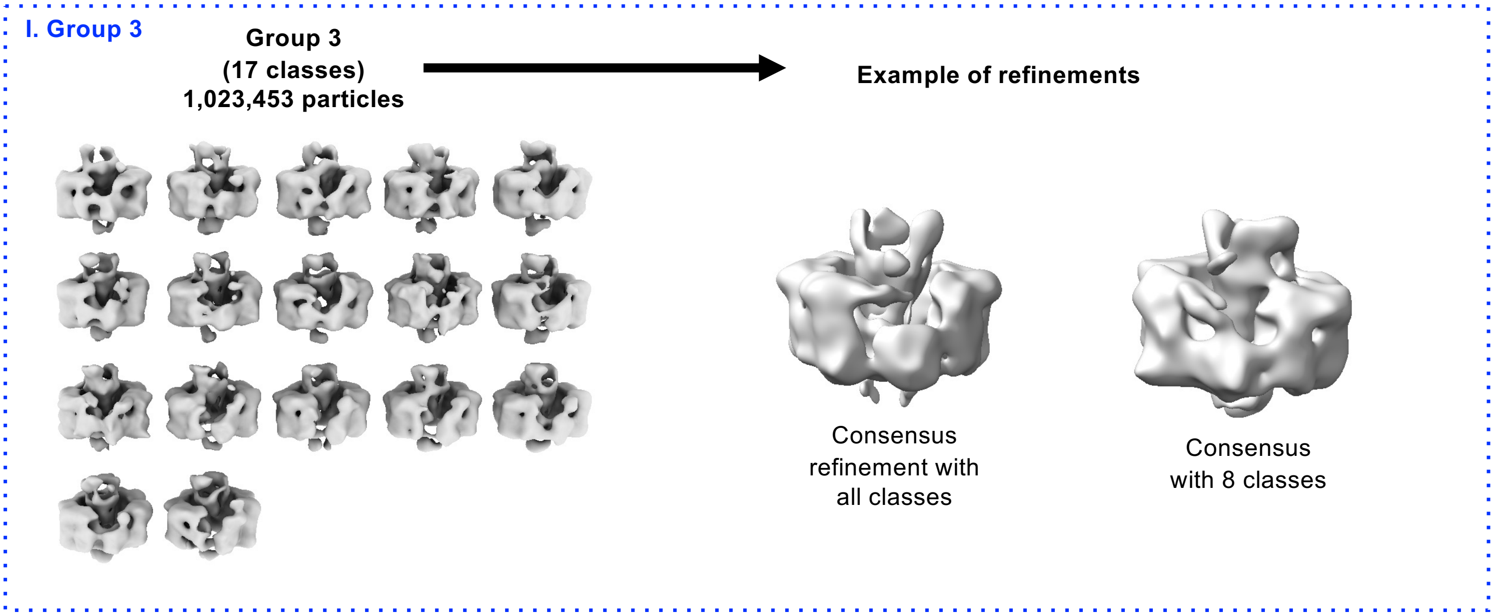


**Figure S6. Cryo-EM data processing workflow for the PA3214 full length structure in complex with PA3213 C-terminal peptide.** *A,* micrographs pre-processing and particle picking. *B,* initial round of cleaning WARP-picked particles that led to the first model. *C,* picking with Topaz and sequential cleaning steps. *D,* classification and refinement of the PA3214 lipoprotein. *E,* refinement and classification of particles leading to full-length PA3214 in closed and in open conformations. *F,* refinement of the full complex. *G,* refinement of the IM region of the complex. *H,* focused refinement of PA3214 from the full consensus map to confirm its orientation relative to the rest of the complex. *I,* 3D reconstructions from particles not included in Map 1 (closed conformation) or Map 2 (open conformation).

#### Supplementary Figure S7

##### A PA3211-PA3214 consensus map

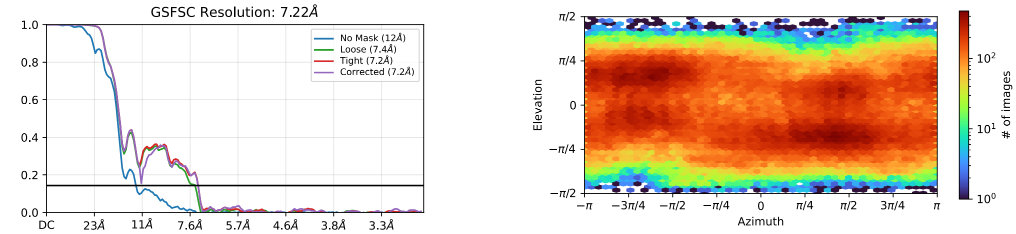


##### B Focused map of PA3214 closed conformation

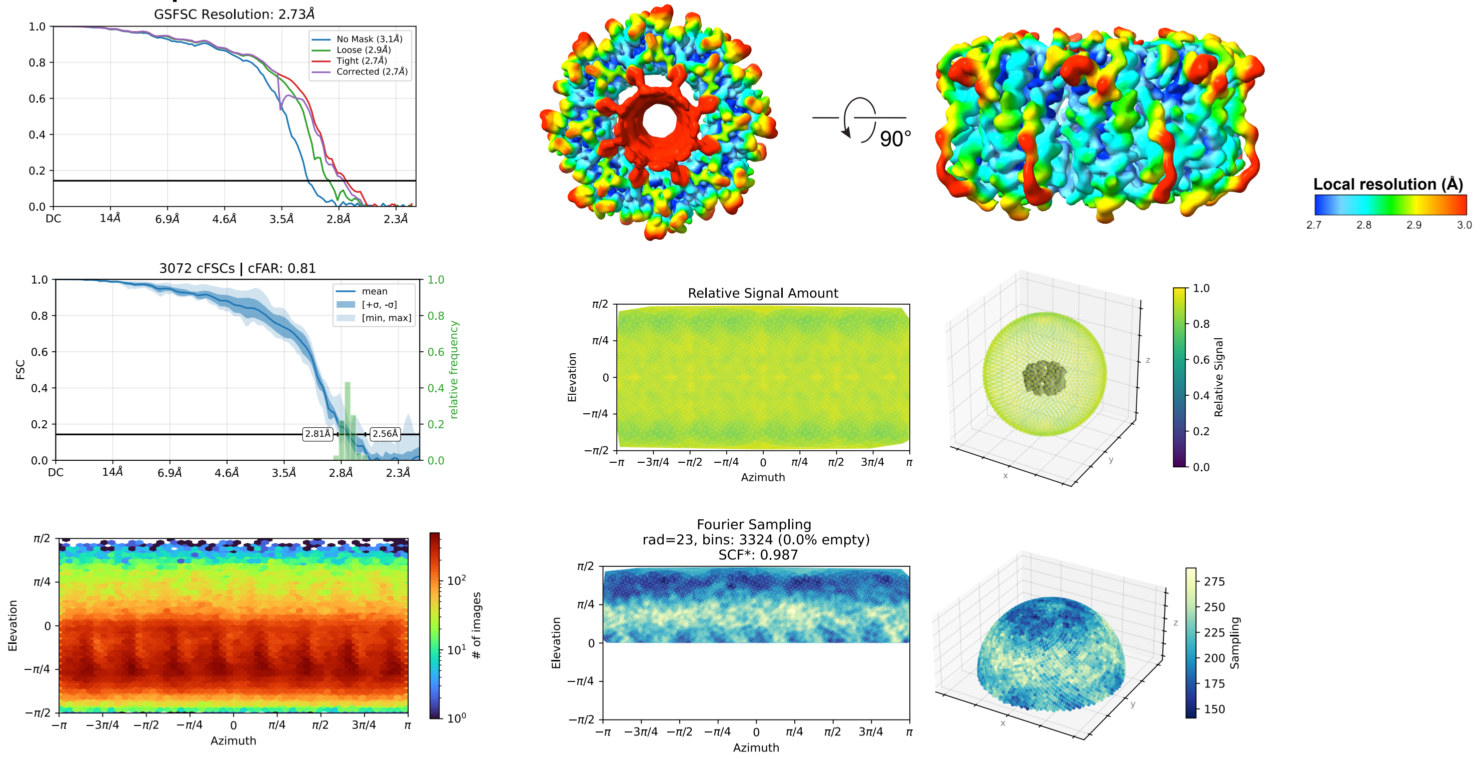


##### C Focused map of PA3214 open conformation

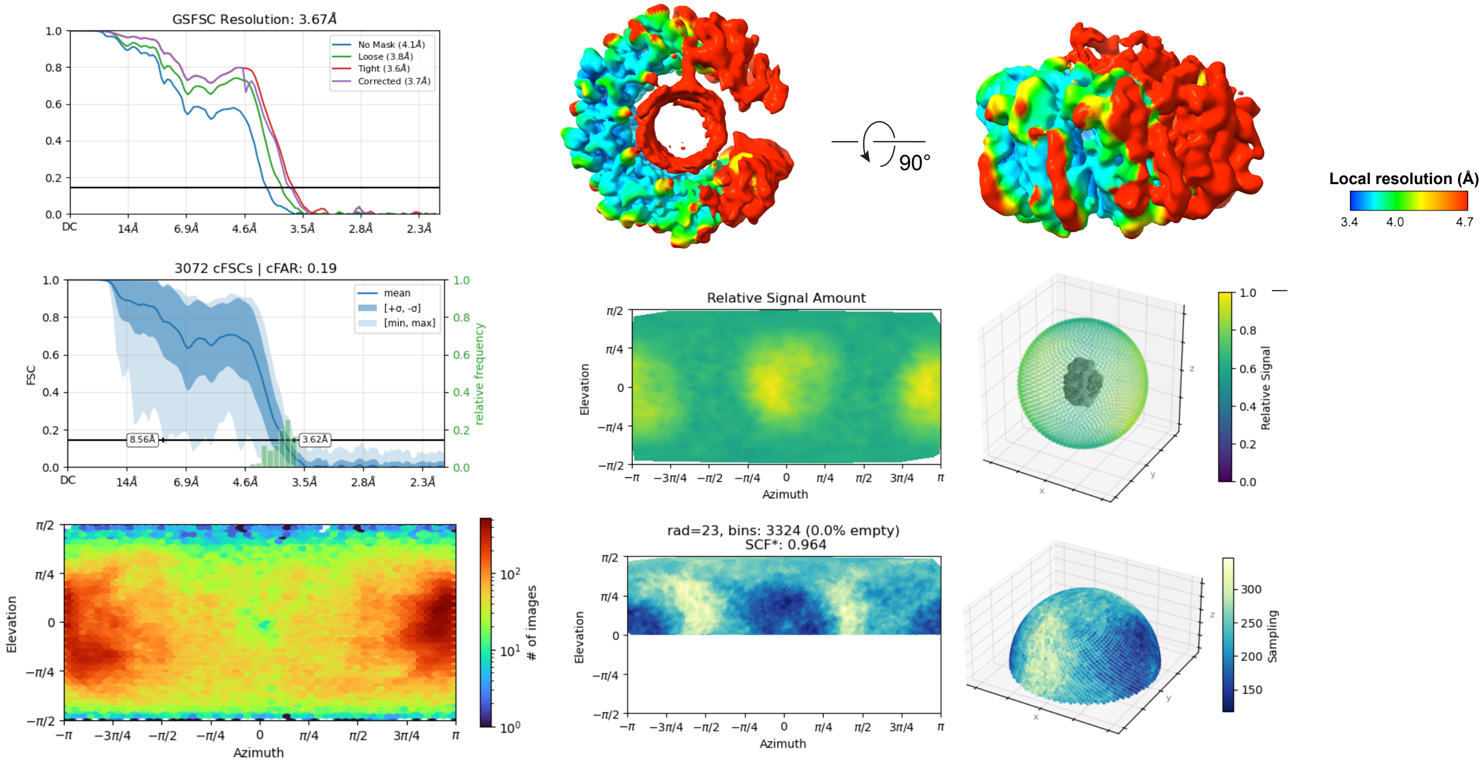

**Figure S7. Map validation corresponding to the workflow in Fig. S6.** *A,* overall resolution and viewing direction distribution of particles in the full consensus map (Map 4). *B,* analysis of the full-length PA3214 map in the closed conformation (Map 2) with the final overall and local resolution (top row), and orientation diagnostics with the cFSC curve, the canonical FSC summary plot, the relative signal amount vs viewing direction (middle row), the viewing direction distribution, and the Fourier sampling plot (bottom row). *C,* analysis of the full-length PA3214 map in the open conformation (Map 3) with the final overall and local resolution (top row), and orientation diagnostics with the cFSC curve, the canonical FSC summary plot, the relative signal amount vs viewing direction (middle row), the viewing direction distribution, and the Fourier sampling plot (bottom row).

Suppplemental Figure S8

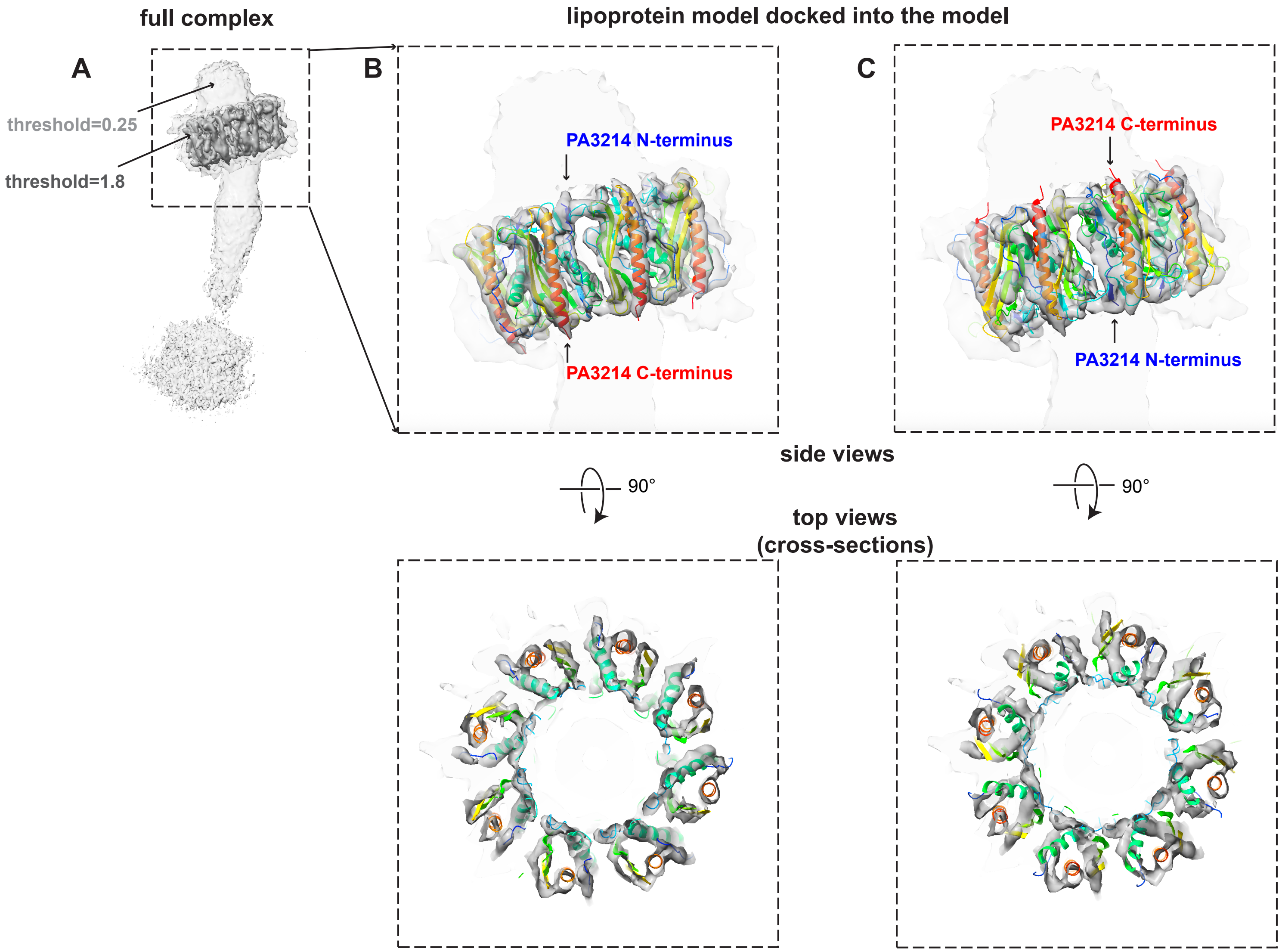


**Figure S8. Local refinement on the consensus map to assign the orientation of PA3214 relative to the rest of the complex.** *A*, locally refined map shown with 2 thresholds, showing the high resolution features of PA3214 as well as the relative position of the PA3211-PA3212-PA3213 domains. *B*, side and top views of the PA3214 model docked into the map with the PA3214 C-termini facing the OM, which accounts for most of the density. *C,* side and top views of the PA3214 model docked into the map with the PA3214 C-termini facing the periplasm, which does not fit well into the density, with regions of the model unaccounted for by density, and other regions with empty density.

**Supplementary Figure S9**

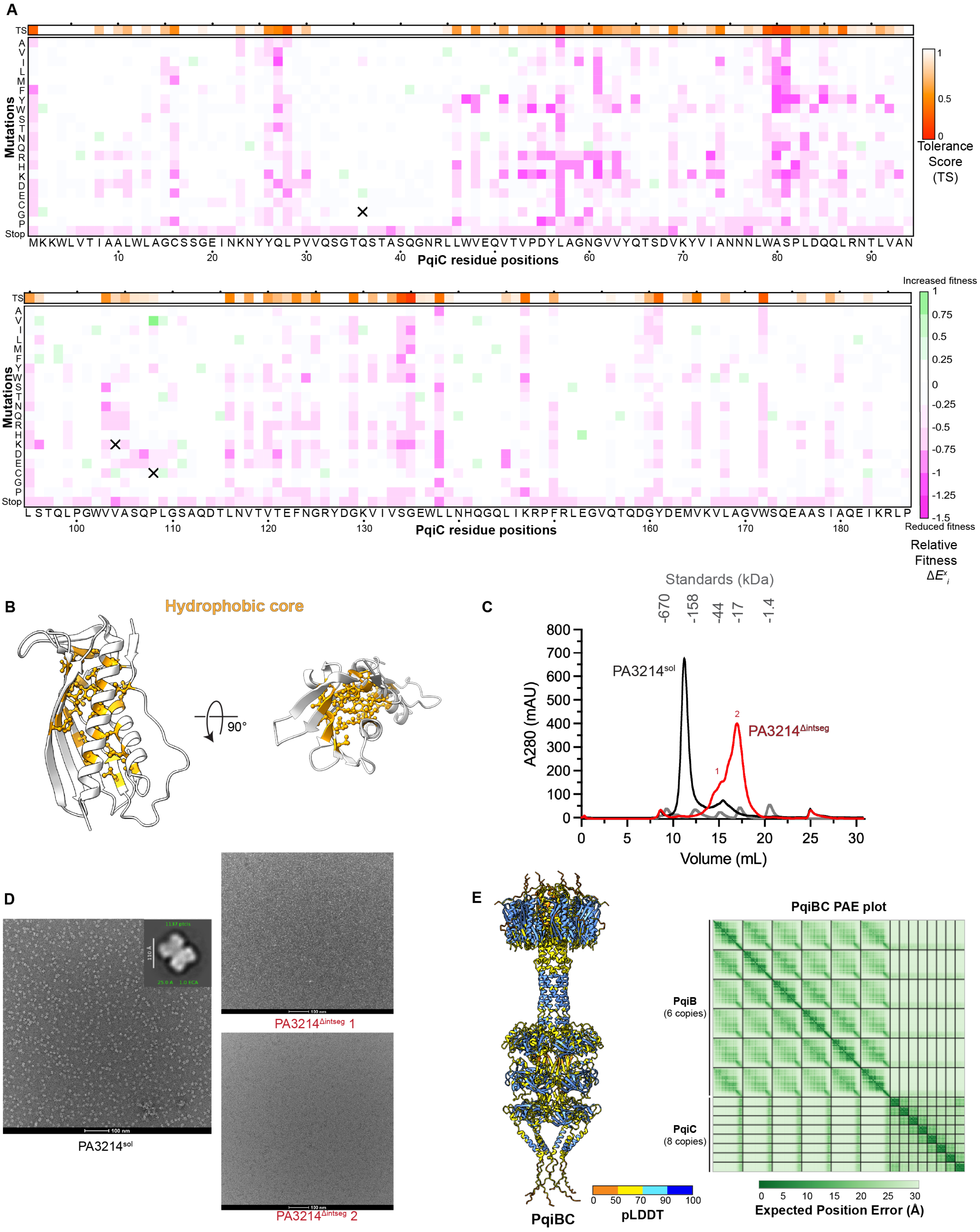

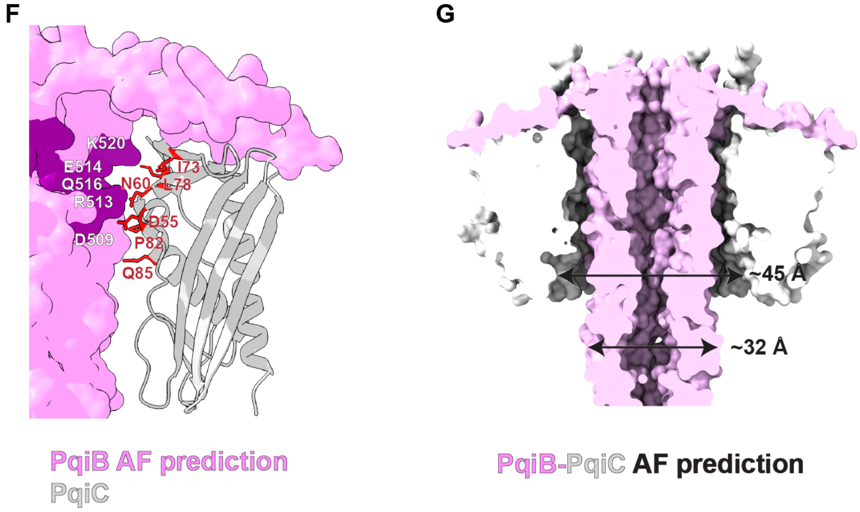


**Figure S9. Results and analysis of PqiC DMS experiments, related to Figures 3-5.** *A,* heat map summarizing the results of PqiC deep mutational scanning (DMS). Each square represents the average fitness cost (over two biological replicates) of an individual mutation relative to the WT sequence. Mutations that are more harmful to fitness relative to the WT are shown in shades of magenta, while mutations that are more beneficial to fitness are in shades of green, and white represents neutral mutations, as shown in the key. The Tolerance Score (calculated as described in the Methods) is shown as the colored square above each strip. Squares that contain an "X" indicate incomplete coverage at that position. *B,* PqiC monomer highlighting the positions of residues with tolerance score <0.9 within the hydrophobic core of the protein (orange). *C,* size exclusion chromatogram for PA3214$^{sol}$ (black) overlaid with PA3214$^{\Delta intseg}$ (red) and standards (gray), with two peaks of PA3214$^{\Delta intseg}$ chosen for analysis by negative staining shown in red numbers. PA3214$^{\Delta intseg}$ elutes between the 44 and 17 kDa protein standards. *D,* representative negative stain micrographs from PA3214$^{sol}$ and PA3214$^{\Delta intseg}$, with an inset of PA3214$^{sol}$ 2D class average, showing oligomers as expected. Neither peak for PA3214$^{\Delta intseg}$ showed clear oligomeric assembly. As PA3214 monomers are only ~20 kDa, we do not expect to resolve these clearly by negative staining. *E,* AlphaFold 3 prediction of PqiBC colored by pLDDT as in the corresponding key, and plot of predicted alignment error (PAE) plotted as a heatmap. *F,* Alphafold 3 PqiB prediction (pink) docked on PqiC (PDB 82QC) (white). The AlphaFold 3 model of the PqiBC complex was predicted, and aligned on the experimental coordinates of PqiC (PDB 82QC) to generate this figure. Potential interactions occurring within the PqiC pore are mapped. *G,* surface representation of PqiBC prediction used in *E*. Black arrows represent the predicted width of the PqiB needle and the PqiC pore.

**Suppplemental Figure S10**

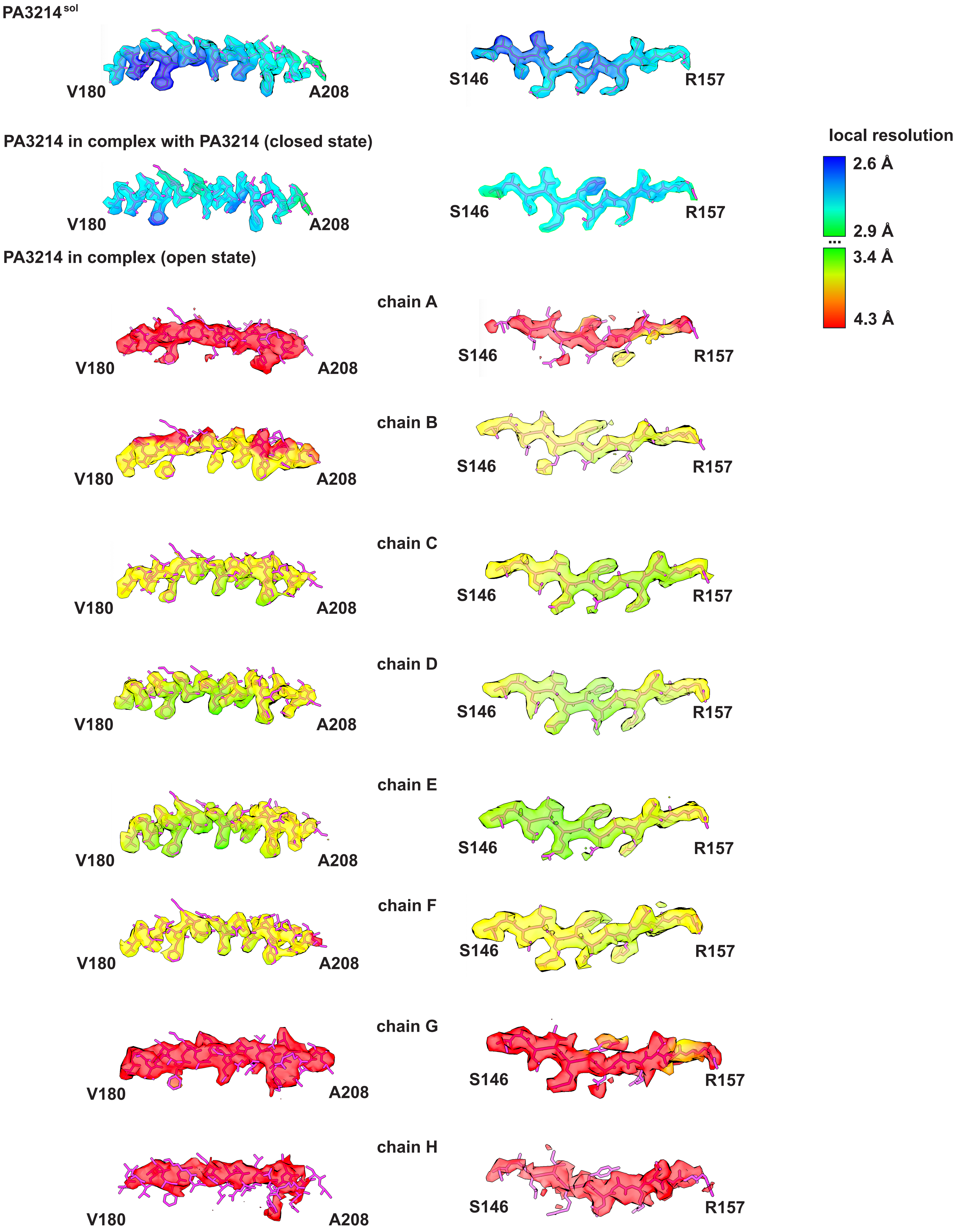

**Figure S10. Representative cryo-EM map densities.** Model to map fits and representative density from a helix (V180-A208) and a beta-strand (S146-R157) from the different PA3214 structures. All maps are colored by local resolution.
