## Supplementary Tables (1-3) for "Interactions of outer membrane lipoproteins *P. aeruginosa* PA3214 and *E. coli* PqiC with their MCE protein binding partners, PA3213 and PqiB"

**Supplementary Table S1A. cryo-EM data collection for PA3214$^{sol}$**

| **Data collection** | |
|---|---|
| Microscope | Krios1 (PNCC) |
| Voltage | 300 |
| Energy filter | None |
| Nominal defocus range (µm) | 0.8-2.1 |
| Camera | Gatan K3 |
| Detector mode | super resolution |
| Nominal pixel size (Å/pixel) | 0.514425 |
| Nominal magnification | 22500x |
| Dose rate (e-/Å$^2$ per frame) | 50 |
| Frame rate (ms) | 48 |
| Exposure time (s) | 2.4 |
| Number of movie frames (no.) | 50 |
| Automation software | SerialEM |
| No, of micrographs at 0° tilt | 6,463 |

**Supplementary Table S1B. Refinement and validation statistics for PA3214$^{sol}$**

| **Data processing** | **Map 1** |
|---|---|
| Number of particles | 654,956 |
| Resolution (Å, FSC=0.143) | 2.67 |
| Map sharpening B-factors (Å$^2$) | -156.2 |
| cFAR | 0.73 |
| SCF | 0.815 |
| Symmetry | C8 |
| Box size (px) | 150 |
| **Model refinement** | |
| Initial model | homology model using Phyre2 |
| Model composition | |
| chains | 8 |
| non-hydrogen atoms | 10784 |
| protein residues | 1384 |
| ADP mean B-factors | 133.29 |
| R.M.S. deviations | |
| rmsd (bonds) | 0.002 |
| rmsd (angles) | 0.564 |
| Molprobity score | 1.25 |
| Clashcore, all atoms | 4.79 |
| Rotamer outliers (%) | 0.00 |
| CaBLAM outliers (%) | 1.21 |
| CA Geometry outliers (%) | 0.61 |
| FSC (model-map; Å, FSC=0.143/0.5) | 2.7 / 2.9 |
| Ramachandran plot (%) | |
| Favored | 98.82 |
| Allowed | 1.18 |
| Outliers | 0.00 |
| Rama-Z | |
| whole | -0.79 |

| | |
|---|---|
| helix | 0.52 |
| sheet | -1.29 |
| loop | -0.54 |
| Map CC (mask) | 0.87 |
| Map CC (box) | 0.85 |
| Map CC (peaks) | 0.73 |
| Map CC (volume) | 0.87 |
| EMD ID | EMD-77071 |
| PDB ID | 13HQ |

**Supplementary Table S2A. cryo-EM data collection for PA3211-PA3214 full complex**

| **Data collection** | |
|---|---|
| Microscope | FEI Titan Krios1 (NYSBC) |
| Voltage | 300 |
| Energy filter | Yes |
| Nominal defocus range (μm) | 0.8-3.5 |
| Camera | Gatan K3 |
| Detector mode | counting |
| Nominal pixel size (Å/pixel) | 1.083 |
| Nominal magnification | 81000x |
| Dose rate (e-/Å$^2$ per frame) | 51.51 (untilted) and 51.43 (tilted) |
| Frame rate (ms) | 200 |
| Exposure time (s) | 6 |
| Number of movie frames (no.) | 30 |
| Automation software | Leginon |
| No, of micrographs at 0° tilt | 6,057 |
| No, of micrographs at 30° tilt | 3,929 |

**Supplementary Table S2B. Refinement and validation statistics for PA3211-PA3214 full complex**

| **Data processing** | **PA3214 in closed conformation (Map 2)** | **PA3214 in open conformation (Map 3)** | **Full map (Map 4)** |
|---|---|---|---|
| Number of particles | 378,252 | 187,311 | 326,143 |
| Resolution (Å, FSC=0.143); (Symmetry) | 3.0 (C1)<br>2.7 (C8) | 3.7 (C1) | 12 (C1) |
| Map sharpening B-factors (Å$^2$) | -162.7 | -181.6 | None |
| cFAR | 0.81 | 0.20 | 0.22 |
| SCF | 0.987 | 0.964 | |
| Box size (px) | 256 | 256 | 396 |
| Pixel size (Å) | 1.083 | 1.083 | 1.444 |
| **Model refinement** | | | |
| Initial model | Alphafold & Modelangelo | closed conformation | |
| Model composition | | | |
| chains | 16 | 8 | |
| non-hydrogen atoms | 11520 | 11040 | |
| protein residues | 1480 | 1424 | |
| ADP mean B-factors | 141.46 | 222.46 | |
| R.M.S. deviations | | | |
| rmsd (bonds) | 0.003 | 0.004 | |
| rmsd (angles) | 0.593 | 0.669 | |
| Molprobity score | 0.91 | 1.05 | |
| Clashcore, all atoms | 1.35 | 2.68 | |
| Rotamer outliers (%) | 0.00 | 0.00 | |
| CaBLAM outliers (%) | 1.69 | 1.72 | |
| CA Geometry outliers (%) | 1.13 | 0.93 | |
| FSC (model-map; Å, FSC=0.143/0.5) | 2.7 / 2.9 | 3.6 / 3.9 | |
| Ramachandran plot (%) | | | |

| | | | |
|---|---|---|---|
| Favored | 97.79 | 98.01 | |
| Allowed | 2.21 | 1.99 | |
| Outliers | 0.00 | 0.00 | |
| Rama-Z | | | |
| whole | -0.37 | -0.73 | |
| helix | 0.27 | -0.60 | |
| sheet | 0.06 | 0.20 | |
| loop | -0.67 | -0.72 | |
| Map CC (mask) | 0.89 | 0.80 | |
| Map CC (box) | 0.88 | 0.86 | |
| Map CC (peaks) | 0.78 | 0.68 | |
| Map CC (volume) | 0.89 | 0.80 | |
| EMD ID | EMD-77072 | EMD-77073 | EMD-77074 |
| PDB ID | 13HR | 13HS | |

**Supplementary Table 3. Bacterial strains and constructs used in this study**

| Reagent/ Resource | Description | Reference | Addgene ID |
|---|---|---|---|
| Rosetta 2(DE3) | *E. coli* competent cells for protein expression (#714003) | Novagen | |
| TOP10 | *E. coli* competent cells for plasmid propagation (#C404010) | ThermoFisher | |
| BW25113 | *E. coli* K-12 parent strain | | |
| bBEL455 | *E. coli* K-12 BW25113 *ΔpqiC* | | |
| pBEL1163 | [6xHis-TEV-PqiB(39-546)] expression vector | This study | 253832 |
| pBEL1282 | PqiA-[6xHis2xQH-TEV-PqiB]-PqiC expression vector | This study | 253833 |
| pBEL1296 | [PelB-6xHis-TEV-PqiC(17-187)] expression vector | This study | 253834 |
| pBEL2277 | [6xHis2xQH-TEV-PA3214(27-214)] expression vector | This study | 253835 |
| pBEL3076 | [6xHis2xQH-TEV-PA3211]-PA3212-PA3213-PA3214 expression vector | This study | 253836 |
| pBEL3081 | PA3211-PA3212-PA3213-[PA3214-2xHQ6xHis] expression vector | This study | 253837 |
| pBEL3085 | pET17b-pqiC for complementation | This study | 253838 |
| pBEL3438 | PqiA-PqiB-PqiC expression vector | This study | 253839 |
| pBEL3439 | PqiA-[6xHis2xQH-TEV-PqiB(1-530)]-PqiC expression vector | This study | 253840 |
| pBEL3440 | PqiA-[6xHis2xQH-TEV-PqiB(D538G)]-PqiC expression vector | This study | 253841 |
| pBEL3441 | PqiA-[6xHis2xQH-TEV-PqiB(P539G)]-PqiC expression vector | This study | 253842 |
| pBEL3442 | PqiA-[6xHis2xQH-TEV-PqiB(P541G)]-PqiC expression vector | This study | 253843 |
| pBEL3444 | PqiA-[6xHis2xQH-TEV-PqiB]-PqiC(Y64K) expression vector | This study | 253844 |
| pBEL3445 | PqiA-[6xHis2xQH-TEV-PqiB]-PqiC(Y161K) expression vector | This study | 253845 |
| pBEL3446 | PA3211-PA3212-PA3213-PA3214 expression vector | This study | 253846 |
| pBEL3447 | PA3211-PA3212-PA3213(1-295)-[PA3214-6xHis2xQH] expression vector | This study | 253847 |
| pBEL3448 | [6xHis2xQH-TEV-PA3214(55-214)] expression vector | This study | 253848 |
| pBEL3449 | PA3211-PA3212-PA3213(E309G)-[PA3214-6xHis2xQH] expression vector | This study | 253849 |
| pBEL3450 | PA3211-PA3212-PA3213(F310G)-[PA3214-6xHis2xQH] expression vector | This study | 253850 |
| pBEL3451 | PA3211-PA3212-PA3213(P312G)-[PA3214-6xHis2xQH] expression vector | This study | 253851 |
| pBEL3452 | PA3211-PA3212-PA3213-[PA3214(R73E)-6xHis2xQH] expression vector | This study | 253852 |
| pBEL3453 | PA3211-PA3212-PA3213-[PA3214(V76K)-6xHis2xQH] expression vector | This study | 253853 |
| pBEL3454 | PA3211-PA3212-PA3213-[PA3214(Y86K)-6xHis2xQH] expression vector | This study | 253854 |
| pBEL3455 | PA3211-PA3212-PA3213-[PA3214(K178E)-6xHis2xQH] expression vector | This study | 253855 |
| pBEL3456 | PA3211-PA3212-PA3213-[PA3214(V180K)-6xHis2xQH] expression vector | This study | 253856 |
